# Immune sensing of food allergens promotes aversive behaviour

**DOI:** 10.1101/2023.01.19.524823

**Authors:** Esther B. Florsheim, Nathaniel D. Bachtel, Jaime Cullen, Bruna G. C. Lima, Mahdieh Godazgar, Cuiling Zhang, Fernando Carvalho, Gregory Gautier, Pierre Launay, Andrew Wang, Marcelo O. Dietrich, Ruslan Medzhitov

## Abstract

In addition to its canonical function in protecting from pathogens, the immune system can also promote behavioural alterations^1–3^. The scope and mechanisms of behavioural modifications by the immune system are not yet well understood. Using a mouse food allergy model, here we show that allergic sensitization drives antigen-specific behavioural aversion. Allergen ingestion activates brain areas involved in the response to aversive stimuli, including the nucleus of tractus solitarius, parabrachial nucleus, and central amygdala. Food aversion requires IgE antibodies and mast cells but precedes the development of gut allergic inflammation. The ability of allergen-specific IgE and mast cells to promote aversion requires leukotrienes and growth and differentiation factor 15 (GDF15). In addition to allergen-induced aversion, we find that lipopolysaccharide-induced inflammation also resulted in IgE-dependent aversive behaviour. These findings thus point to antigen-specific behavioural modifications that likely evolved to promote niche selection to avoid unfavourable environments.

---

Allergies are a class of inflammatory diseases that have increased in prevalence over recent decades^4^. Allergic diseases such as atopic dermatitis, food allergies, asthma, and drug hypersensitivities seem to be directly linked to industrialization and urban lifestyles^5–7^. The physiological roles for these allergic responses, however, remain enigmatic. Type 2 immunity, which includes Th2 lymphocytes, IgE antibodies, and innate immune cells (e.g., mast cells, eosinophils, and type 2 innate lymphoid cells), mediates allergic responses. When chronic or excessive, allergic responses become detrimental, and potentially lethal^8^. Allergic responses appear to play an important role in host defence against noxious substances, including venoms, hematophagous fluids, xenobiotics, and irritants^9–12^. Indeed, a common feature of allergic responses is the exacerbation of defensive neuronal reflexes like sneezing, itching, and vomiting, which expel harmful substances from the body^13^. In addition to these reflexes, avoidance behaviour was shown to be induced in allergic responses^14–16^, which suggests that type 2 immunity might limit exposure to detrimental stimuli, acting as an efficient defence strategy to prevent further damage. However, the mechanisms by which type 2 responses promote behavioural outputs have yet to be established.

To examine the impact of allergic sensitization on avoidance behaviour, we sensitized mice with subcutaneous injections of ovalbumin (OVA) and the adjuvant aluminium hydroxide (alum) on days 0 and 7 (Fig. 1a). Control mice received alum only. Mice were then acclimatized to home cages equipped with two lickometers (i.e., spouts that automatically detect licks) connected to water bottles. During the acclimation period, mice showed no increased preference for any of the water bottles (Extended Data Fig. 1a, b). We then randomly switched the content of one of the bottles to an OVA solution. Control mice from both BALB/c and C57BL/6 backgrounds showed an increased preference for the OVA solution compared to water (Fig 1b and Extended Data Fig. 1c), suggesting that the OVA solution is appetitive for mice. In contrast, sensitized mice showed a decreased preference to the OVA solution in a dose-dependent manner on the first day of preference test (Fig. 1b and Extended Data Fig 1d, e). This change in OVA preference by sensitized mice occurred as early as 10 min after providing the test bottles (Extended Data Fig. 1f). Notably, aversion to the OVA solution persisted for at least 48 weeks after allergic sensitization (Fig. 1d) and it was specific to OVA, since control and sensitized mice showed comparable preference to a solution containing bovine serum albumin (BSA) (Fig. 1e). We next found that the transient receptor potential cation channel subfamily M member 5 (TRPM5), required for taste transduction in chemosensory cells^17^, was dispensable for the development of allergen aversion (Fig. 1f). Oral sensitization with cholera toxin, a commonly used adjuvant to induce experimental food allergy, also promoted aversion to OVA (Extended Data Fig. 1g). Together, our data indicate that immunization towards a protein can generate specific food aversions, which is consistent with previous observations^14,18^. Since we found that mice naturally preferred OVA alone, we did not add sucrose to the OVA solution, as prior studies did, to minimize behavioural and metabolic effects^19^. Thus, allergic immunization induces long-lasting and antigen-specific avoidance behaviour independent on the protein taste.

**Fig. 1.**
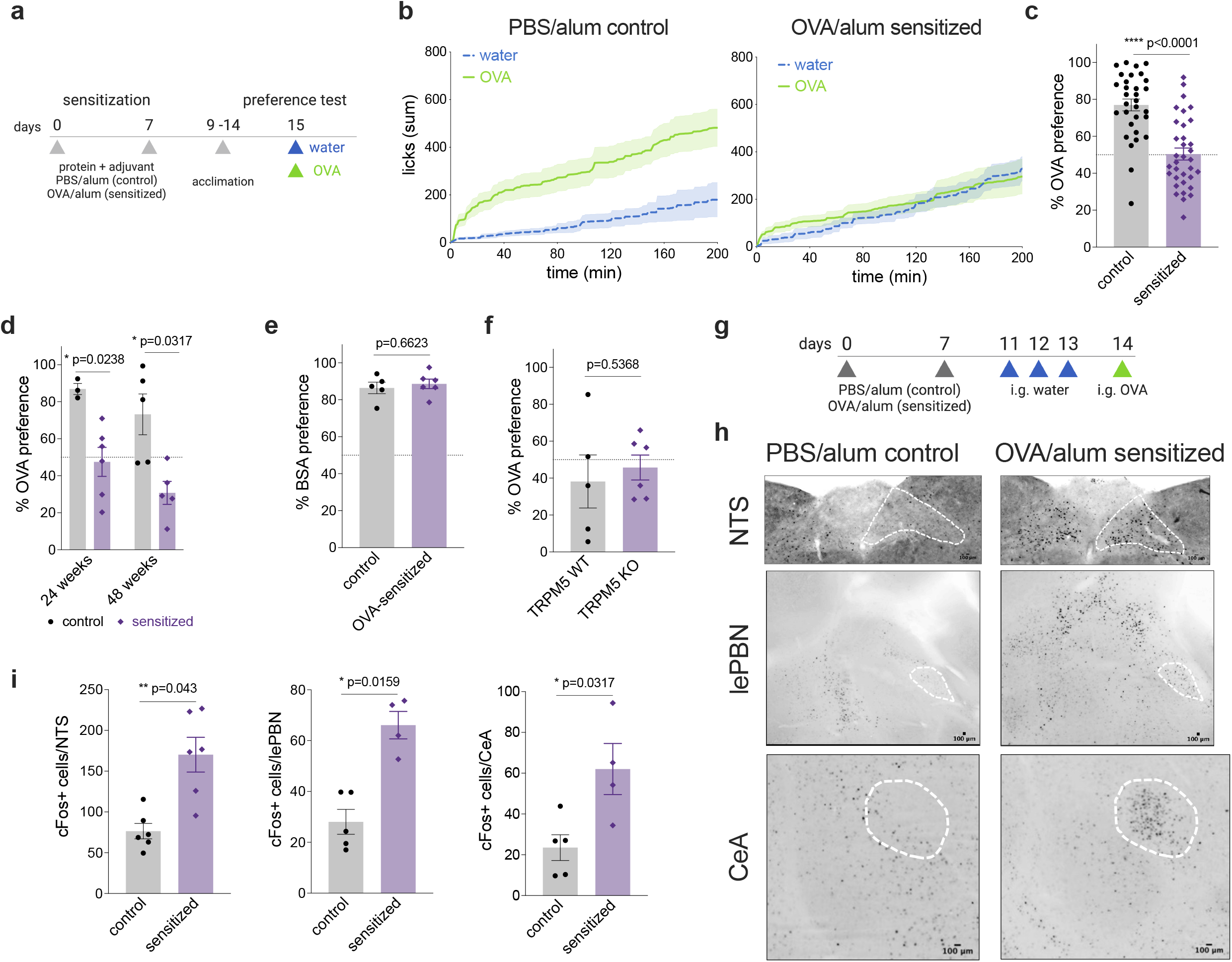
Allergic sensitization induces specific and long-lasting aversion to food allergen. **a**, Schematic protocol for allergic sensitization and behavioral assay. **b**, Cumulative licks from mice sensitized with PBS/alum or OVA/alum. Preference test consisting of one water bottle and one 1% OVA bottle on day 1. **c**, Preference to OVA solution. **d**, Preference to OVA after 24 or 48 weeks of sensitization with alum or OVA/alum. **e**, Preference to bovine serum albumin. **f**, Preference to OVA by OVA/alum sensitized TRPM5 WT or KO mice. **g**, Schematic protocol of allergic sensitization and oral challenge. Mice were orally challenged with OVA after sensitization and 3 sham gavages with water. Controls received all but OVA during sensitization. **h**, Immunofluorescence images of the *nucleus tractus solitarius* (NTS, top), lateral external parabrachial nucleus (lePBN, middle), and central amygdala (CeA, bottom) from control (n=5) or OVA/alum sensitized (n=4) BALB/c mice using anti-cFos antibody, 90 min after one OVA challenge. Scale bars = 100 µm. **i**, Number of cFos+ neurons in the NTS, lePBN, and CeA of BALB/c control (n=5) or OVA/alum (n=4) sensitized mice. Graphs show mean ± s.e.m., Mann-Whitney test. Representative of at least 2 experiments.

Aversive responses to unpleasant stimuli were previously shown to induce brain activation within the nucleus of tractus solitarius (NTS), lateral external parabrachial nucleus (lePBN), and central amygdala (CeA)^20–22^. To determine whether the ingestion of allergens can activate these brain areas, we orally challenged control and sensitized mice with OVA and 90 minutes later collected their brains to test for neuronal activation using cFos as a marker (Fig. 1g). We found that one allergen challenge was enough to induce NTS, lePBN, and CeA activation in sensitized mice as compared with controls (Fig. 1h, i). There was no difference in cFos staining of the lateral hypothalamus or the paraventricular nucleus of the hypothalamus (LH and PVN, respectively; Extended Data Fig. 1h, i). Our results demonstrate that sensitization with OVA leads to prototypical neuronal activation in the central nervous system. These brain regions correlate with aversive behaviour toward the sensitized protein and are likely triggered as a defence for limiting allergen intake.

Increased antigen-specific IgE is a hallmark of allergic sensitization and widely used for clinical diagnosis of allergies^23,24^. As total and OVA-specific IgE antibodies are increased in the circulation two weeks after the first allergen sensitization (Fig. 2a), we examined whether these antibodies could mitigate the avoidance to allergen using IgE-deficient mice. The sensitization to OVA in IgE KO mice did not result in decreased preference to OVA solution compared with their sensitized littermate WT controls (Fig. 2b). Instead, IgE KO mice continued preferring the OVA solution over water (Fig. 2c). Consistently, sensitized mice that are deficient in the high affinity receptor for IgE, FcεRI, showed similar preference to OVA as the WT unsensitized controls (Extended Data Fig. 2a). As expected, production of IgE in sensitized FcεRI KO was comparable to that of WT mice (Extended Data Fig. 2b). By using chimeric mice, we found that aversion to OVA was also dependent on hematopoietic cells expressing FcεRI (Fig. 2d). The increased OVA preference observed in sensitized WT mice reconstituted with FcεRI KO hematopoietic cells (Fig. 2e) was likely due to downstream effects of IgE signalling as the serum levels of OVA-specific IgE were comparable with the WT mice reconstituted with WT hematopoietic cells (Fig. 2f).

**Fig. 2.**
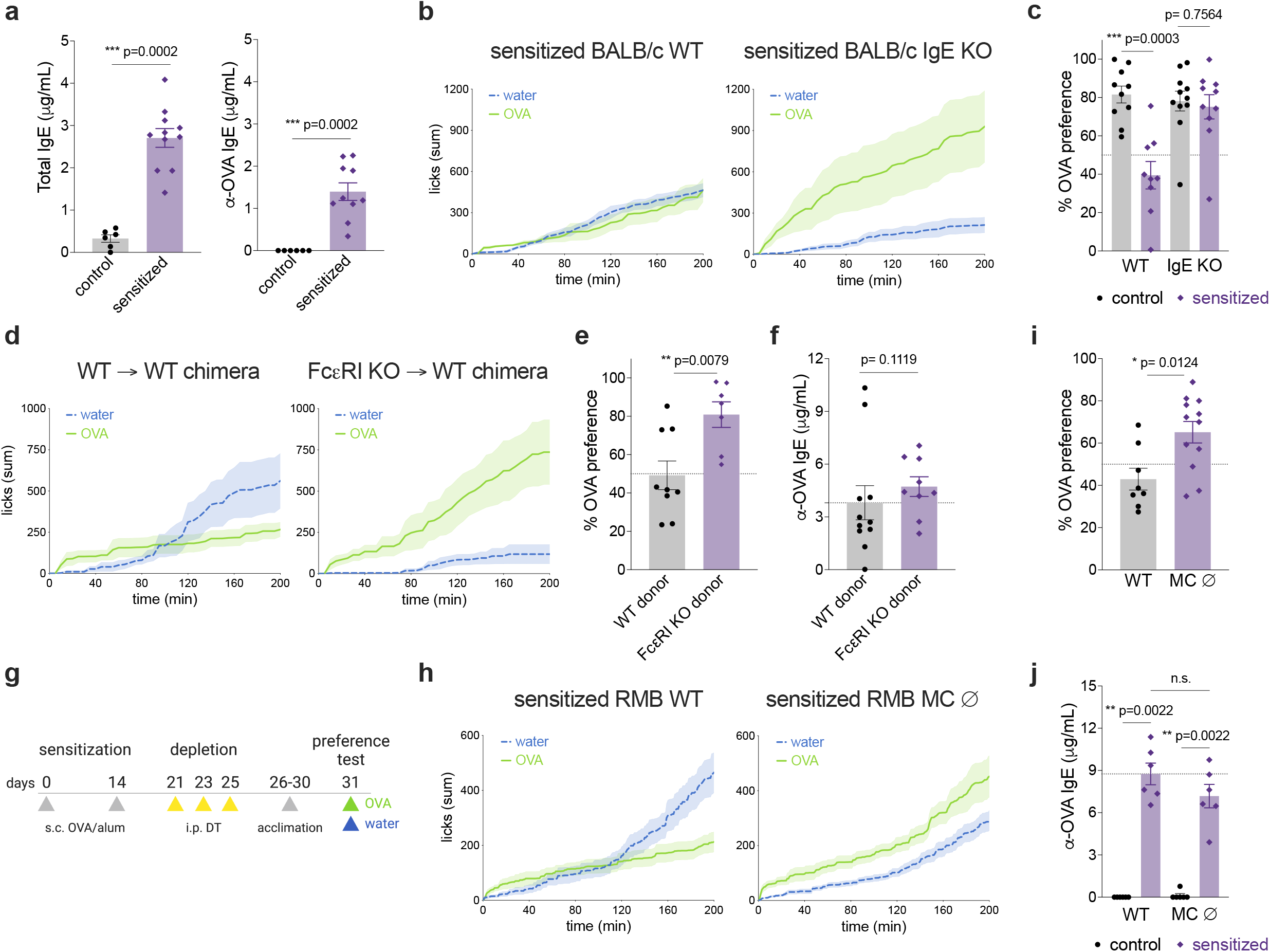
Allergic aversion requires IgE and mast cells. **a**, Total (left panel) and OVA-specific (right) levels of serum IgE on day 14 after allergic sensitization in BALB/c mice. **b**, Cumulative licks to water and OVA solutions during the two-bottle preference test in OVA/alum sensitized BALB/c IgE KO (right) and WT littermate controls (left). **c**, Drinking preference to OVA bottles in controls and allergic sensitized BALB/c WT or IgE KO mice. **d**, Cumulative licks to water and OVA bottles in OVA/alum sensitized C57BL/6 WT or FcεRI chimeras. C57BL/6 WT or FcεRI KO bone marrow hematopoietic cells were transplanted into irradiated WT recipients. **e**-**f**, Drinking preference to OVA bottles (**e**) and OVA-specific IgE serum levels (**f**) in allergic sensitized C57BL/6 WT or FcεRI chimeras. **g**, Schematic protocol of FcεRI+ cell depletion with diphtheria toxin in RMB BALB/c mice. **h**, Cumulative licks to water and OVA bottles in allergic sensitized and DT-injected RMB WT (left) and RMB mutants (right). **i**-**j**, Preference to OVA solution (**i**) and OVA-specific IgE (**j**) in DT-injected RMB WT and mutants. Graphs show mean ± s.e.m, Mann-Whitney test. Representative of at least 2 experiments.

Because IL-4 is required for IgE production and the development of type 2 immune responses^25,26^, we next tested the role of IL-4 signalling through IL-4Ra in the induction of avoidance behaviour to OVA. We used a one-bottle assay for seven days to quantify the consumption of OVA solution when water was not provided. We found that OVA/alum sensitized IL-4Ra KO mice did not decrease their consumption of OVA (Extended Data Fig. 2c), suggesting a role for IL-4 signalling in the development of food aversion. Eosinophils are duodenum-resident cells that have previously been implicated in the initiation of type 2 responses^27^. To determine the role of eosinophils in avoidance behaviour, we used eosinophil-deficient mice, Gata1Δ. These mice showed decreased consumption of OVA when sensitized (Extended Data Fig. 2c), suggesting that eosinophils might be dispensable for induction of allergen aversion. We found that sensitized Gata1Δ, but not IL-4Ra KO, responded to systemic allergen challenge by inducing anaphylaxis (Extended Data Fig. 2d) and showed increased levels of IgE antibodies (Extended Data Fig. 2e). Interestingly, the IgE-dependent induction of allergen aversion was not limited to allergic sensitization, as mice sensitized with OVA and lipopolysaccharide (LPS) – an established non-allergic inflammatory stimulus – also showed decreased preference to OVA, which required IgE (Extended Data Fig. 2f, g). Indeed, OVA/LPS sensitized WT mice had low but increased levels of IgE (Extended Data Fig. 2h), which might be sufficient to promote avoidance to a conditioned antigen. Finally, the lack of avoidance behaviour in OVA/alum sensitized IgE-deficient mice was not due to an intrinsic inability to induce aversion because they showed strong aversion to a bitter compound, denatonium benzoate (DB) (Extended Data Fig. 2i). These data show that avoidance behaviour induced by both allergic and non-allergic immunological stimuli depends on IgE antibodies.

Our findings so far point to a protective role for IgE antibodies. However, these antibodies are known to be detrimental and drive allergic symptoms upon exposure to food allergens^28,29^. We hypothesize that, at early stages, IgE promotes allergic defences such as avoidance behaviour, but upon chronic exposure to allergens, IgE leads to disease^30^. To better define the role of IgE on gut allergic inflammation, we orally challenged mice with OVA 5 times after sensitization as an experimental model of food allergy (Extended Data Fig. 3a). We challenged mice systemically on day 14 to induce anaphylaxis. We found that IgE antibodies are not required for the induction of systemic anaphylaxis (Extended Data Fig. 3b), as reported^31^. This is likely due to OVA-specific IgG1 antibodies that are increased by allergic sensitization in an IgE-independent way (Extended Data Fig. 3c). As previously shown^32^, we found that IgE is essential to accelerate gut motility (Extended Data Fig. 3d) and diarrhoea (data not shown) in response to acute allergen exposure in the gastrointestinal tract. In the intestinal tissue, the accumulation of mast cells in the epithelial and lamina propria compartments, as well as lamina propria eosinophils, was partially dependent on IgE in sensitized and orally challenged mice (Extended Data Fig. 3f). These data demonstrate that IgE endorses pathological processes upon chronic stimulation with allergen, promoting increased GI peristalsis and inflammatory cellular infiltrates in the allergic small intestines.

Mast cells are major gut-resident immune cells implicated in allergies, anaphylaxis, inflammatory bowel disease, and abdominal pain^32–34^. To address the role of mast cells in the development of avoidance behaviour to allergens, we sensitized RMB mice with OVA/alum. We depleted mast cells in these mice with 3 injections of diphtheria toxin, as previously described^35^ prior to the preference test (Fig. 2g). Despite being sensitized with OVA/alum, mast cell depleted-RMB mice showed higher consumption of OVA solution (Fig. 2h) and OVA preference (Fig. 2i) compared with control mice. The systemic levels of IgE antibodies were equally increased in mast cell-depleted mice compared with sensitized WT (Fig. 2j), suggesting that mast cells can drive avoidance through IgE sensing of allergens.

Diverse mast-cell derived mediators have been implicated in neuronal excitation in the gastrointestinal tract during intestinal anaphylaxis^13,36^. Given the rapid change of behaviour observed, we hypothesized that the responsible mediators must either be pre-formed or synthesized de novo after IgE-dependent cross-linking^37^. Histamine and serotonin, released upon mast cell degranulation, are two pre-formed mediators with known roles in mediating itch, pain, diarrhoea, and visceral malaise^32,38–42^. We tested the role of the histamine receptors H1 and H2 using the inhibitors loratadine and famotidine. Acute pre-treatment with both drugs did not affect OVA preference in control or allergic sensitized mice (Fig. 3a), suggesting that histamine might not contribute to aversion to food allergens. Similarly, blockade of serotonin synthesis through 5 days of pre-treatment with parachlorophenylalanine led to only a mild and variable effect on the aversion (Fig. 3a). We found no effect of pre-treatment with the serotonin receptor 5-HT3 antagonist, ondansetron, as compared to vehicle-treated controls (Extended Data Fig. 4a). Mast cell-nociceptor circuits are well described in the skin, lung, and gastrointestinal tract and are proposed to contribute to inflammation and pain perception. Two mediators well-known for these interactions are substance P and CGRP^43,44^, which we tested to determine their possible role in mediating the aversive response. Using pharmacological (substance P receptor inhibitor, aprepitant) and genetic approaches (substance P KO mice), we found that substance P did not affect aversive responses to OVA after allergic sensitization (Extended Data Fig. 4 a, b). We also found that sensitized mice treated with a CGRP receptor inhibitor (BIBN4096) developed aversive behaviour towards OVA comparable to vehicle-treated mice (Extended Data Fig. 4c). Mast cells can release proteases that activate submucosal neurons through cleavage of PAR2^45^, however, pre-treatment with the PAR2 inhibitory peptide FSLLRY-NH2 had no effect on the aversive response of sensitized mice (data not shown). So far, our results indicate that histamine, serotonin, substance P, CGRP, and PAR-2 are not required for aversion to food allergens. It is possible, however, that we were not able to fully suppress the action of some of these mediators due to timing and dosing issues.

**Fig. 3.**
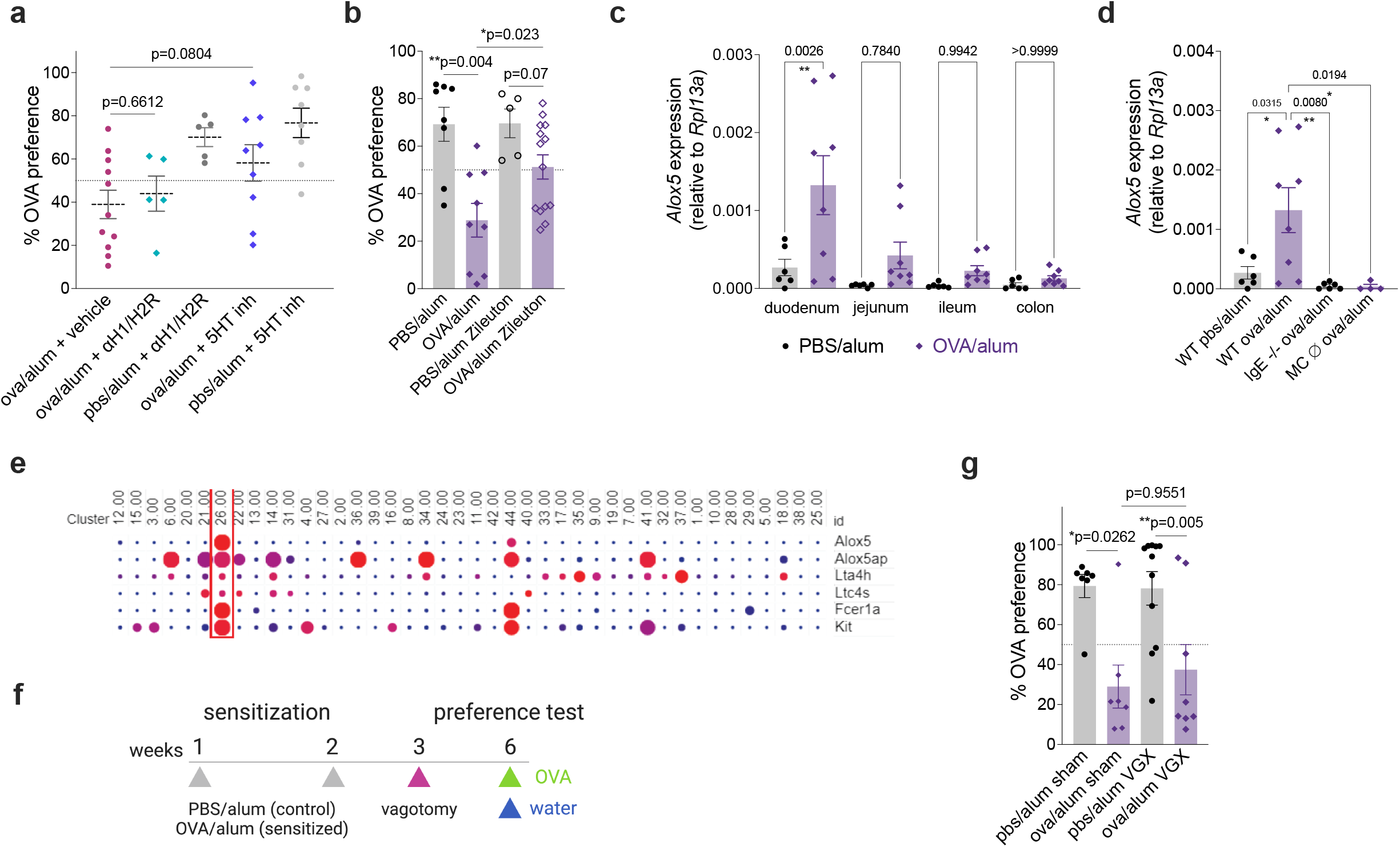
Allergen-induced avoidance behavior requires cysteinyl leukotrienes. **a**, Drinking preference to OVA solution by OVA/alum sensitized mice. Groups were administered with antagonists for H1 and H2 histamine receptors (loratadine and famotidine), a serotonin synthesis inhibitor (parachlorophenylalanine), or a vehicle solution before the two-bottle preference test. n=5-11 per group. **b**, Preference to OVA in OVA/ alum sensitized mice. Zileuton was used as a 5-lipoxygenase inhibitor 1 h prior to the preference test. n=5-14, pooled from 2 experiments. **c**, *Alox5* expression across the GI tract of control and OVA/alum sensitized mice after allergen oral challenges. **d**, *Alox5* expression in the duodenum of OVA sensitized WT, IgE-deficient, or mast cell-depleted mice. **e**, Comparison of gene expression in BALB/c intestinal immune cells from scRNA-seq data of control mice. Cluster 26 represents mast cells whereas cluster 44 refers to basophils. **f**, Schematic protocol for subdiaphragmatic vagotomy in OVA/alum sensitized mice. **g**, OVA preference after vagotomy. **a** - **d, g**, Mean ± s.e.m. **a, b, g**, Mann-Whitney test. **c, d**, One-Way ANOVA with multiple comparisons. **a, c, d, g**, Representative of at least two experiments.

In addition to pre-formed substances, mast cells produce arachidonic acid-derived lipid mediators in minutes after IgE-mediated degranulation^46–48^. Prostaglandins and leukotrienes are known to be produced by mast cells and have profound effects on behaviour through actions on nociceptors and vagal neurons^49–53^. Prostaglandins and leukotrienes are generated by a series of enzymatic steps controlled by rate-limiting cyclooxygenase (COX) and lipoxygenase (LOX) enzymes, respectively. Although COX1/2 inhibition with indomethacin had no effect on the magnitude of the aversive response (Extended Data Fig. 4d), aLOX5 inhibition by pre-treatment with zileuton significantly increased the preference for OVA in sensitized mice, while not impacting OVA preference in controls (Fig. 3b). Using organ homogenates from BALB/c mice sensitized and challenged with intragastric OVA, we determined that *Alox5* expression was transcriptionally induced proximally to distally across the gastrointestinal tract, with the highest induction found in the duodenum (Fig. 3c), and in the epithelium (Extended Data Fig. 4e). This pattern of expression was compared to that of intestinal mast cell expansion (Extended Data Fig. 3e and Extended Data Fig. 6b), and detection of *Alox5* transcripts was largely lost in the duodenum of sensitized IgE KO and mast cell-depleted mice (Fig. 3d). Tuft cells are epithelial cells that can be activated by type 2 inflammation and have the capacity to produce leukotrienes, and this partially depends on TRPM5 signalling^17^. Because TRPM5 KO mice did not display a reduced magnitude of aversion (Fig. 1f), these cells may not be the major source of leukotrienes in this model. Instead, by analysing a previously published single-cell RNA-seq data set of intestinal immune cells in sensitized and challenged BALB/c mice^54^, we found that mast cells and basophils, but not other immune cells, expressed all the transcriptional machinery necessary for leukotriene synthesis (Fig. 3e). Only mast cells appear to express Ltc4s, the enzyme necessary to produce cysteinyl leukotrienes. Taken together, these data suggest that leukotrienes, likely produced by gastrointestinal mast cells, drive avoidance behaviour to food allergens.

Leukotrienes are known to mediate unfavourable sensations through actions on dorsal-root ganglion nociceptors and gut innervating vagal neurons. Indeed, leukotrienes can activate duodenal vagal neurons through Cysltr2 receptors, which are associated with nausea, and nonpeptidergic nociceptors can be active via similar mechanisms leading to itch^40,49^. Following this logic, we probed a role for gut-innervating vagal neurons by conducting subdiaphragmatic vagotomy (Fig. 3f). Successful subdiaphragmatic vagotomy was confirmed by injecting fluorogold intraperitoneally and finding the absence of dye in the dorsal motor nucleus of the vagus (Extended Data Fig. 4f). However, vagotomy had no significant effect on either the preference to OVA in control mice or the development of avoidance in sensitized mice (Fig. 3g). We also sought to examine the role of TRPV1-positive neurons by treating mice with resiniferatoxin (Extended Data Fig. 4g), a potent capsaicin analogue commonly used to ablate these fibres. We confirmed successful ablation by lack of behavioural response and temperature drop to i.p.-injected capsaicin (data not shown). Ablation of these neurons slightly decreased the baseline preference of control mice to OVA (Extended Data Fig. 4h). Because gustatory sensory fibres lack TRPV1 expression^55^, these results suggest that TRPV1+ sensory neurons might control the perceived valence of dietary proteins. Overall, these findings suggest that vagal neuronal afferents individually may not play a dominant role in allergen aversion and that instead mast cell-derived signals such as leukotrienes might be sensed through other pathways.

We next examined whether a humoral pathway is involved in allergen aversion. The area postrema is a sensory circumventricular organ with renowned roles in mediating nausea in the context of noxious stimuli^56^. Growth and differentiation factor 15 (GDF15) is a TGFβ superfamily cytokine produced during conditions of inflammation and cell stress that acts on the area postrema and NTS by binding to its receptor, GFRAL, to mediate conditioned flavour aversion and anorexia^57,58^. Serum GDF15 levels were, in fact, induced upon intragastric food allergen challenge (Fig. 4a), and this induction was amplified with the increasing number of challenges (Extended Data Fig. 5a). Increase in GDF15 depended strongly on the mouse background strain, with C57BL/6 mice displaying no elevation in GDF15 whereas BALB/c displayed a robust elevation of approximately 3-fold after 6 challenges (Extended Data Fig. 5b). This induction was entirely dependent on IgE- and Fcεr1a-expressing cells and could be partially prevented by pre-treating mice with zileuton (aLOX5 inhibitor) prior to each challenge (Fig. 4b, Extended Data Fig. 5c-e), suggesting that mast cell production of leukotrienes may be involved in GDF15 secretion.

**Fig. 4.**
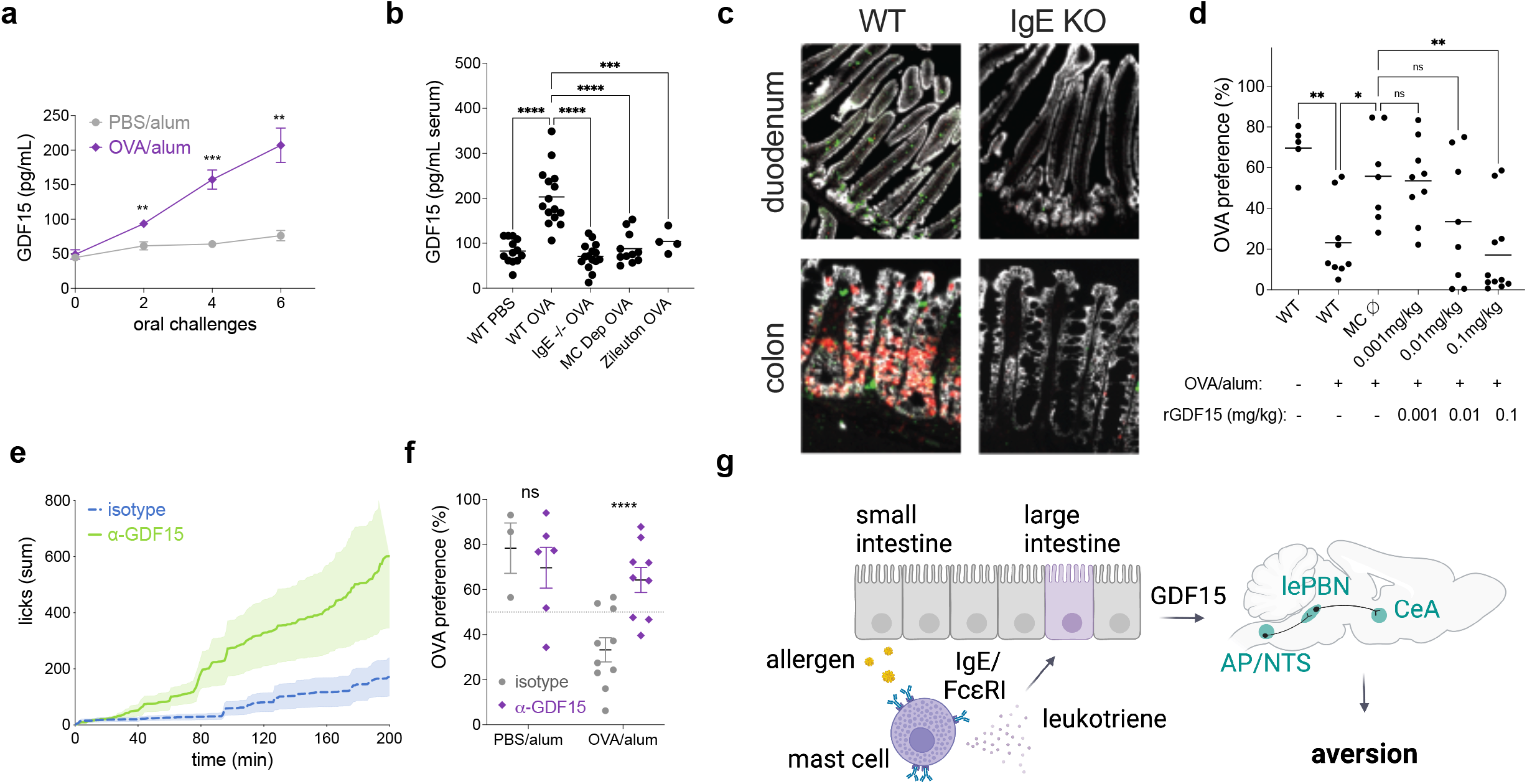
Allergen-induced aversion requires GDF15. **a**, Serum levels of GDF15 in sensitized BALB/c mice after oral allergen challenges. **b**, Serum GDF15 in BALB/c mice after 6 oral challenges with OVA. **c**, Expression of GDF15 mRNA by RNAScope in the duodenum and colon of allergen sensitized and challenged WT or littermates IgE KO mice. **d**, OVA preference 1 h after administration of recombinant mGDF15 in WT and mast cell-depleted (MC ∅) RMB mice. **e**, Cumulative licks on OVA bottle during preference test in OVA/alum sensitized WT mice 5 h after injection with blocking GDF15 antibody or isotype control antibody. **f**, Sensitized WT mice were injected with blocking GDF15 antibody and the OVA preference was quantified 5 h later. **g**, Working hypothesis: allergen is sensed in the gastrointestinal mucosa of sensitized animals through allergen-specific IgE antibodies and FcεRI receptors that are highly expressed by tissue-resident mast cells. Allergen sensing by intestinal mast cells promote the release of cysteinyl leukotrienes, which mediates the secretion of GDF15 by epithelial cells in the large intestine. Systemic GDF15 likely acts directly in the brainstem, activating neurons in the *nucleus of tractus solitarius* (NTS) and area postrema to induce aversive circuits through the parabrachial nucleus (lePBN) and central amygdala (CeA). This regulated organismal response allows for rapid identification of allergens and the development of avoidance behavior, evolved to limit the exposure to potentially noxious stimuli and protect from further damage. **a, b, d, f**, Mean ± s.e.m. **a, b, d** One-Way ANOVA with multiple comparisons. **f**, Mann-Whitney test.

Using organ homogenates from sensitized mice challenged intragastrically with OVA, we found that GDF15 was induced on the transcriptional level mainly in the duodenum and colon (Extended Data Fig. 5f). GDF15 induction in these tissues was dependent on IgE antibodies and mast cells (Extended Data Fig. 5g). Using RNAscope *in situ* hybridization with probes specific for GDF15, Fcεr1a, and EPCAM, we demonstrated that GDF15 was induced by the colon after intragastric OVA challenges, with a lower magnitude of induction in the small intestine (Fig. 4c, Extended Data Fig. 6a-c). This was surprising, as mast cell expansion occurred in a proximal-to-distal pattern across the small and large intestine. In fact, GDF15 transcripts showed little (<1%) overlap with FcεR1a transcripts, and instead colocalized primarily (93%) with EPCAM+ cells (Extended Data Fig. 6d). These GDF15+ epithelial cells were found to form close contact with mast cells and were primarily found in the colonic crypt/transit amplifying zone. Interestingly, similar colonic induction patterns were previously described upon treatment of mice with metformin^59^. Hence, food allergen ingestion induces colonic epithelial GDF15 transcription through IgE and mast cells.

To determine whether GDF15 was able to impact the aversive response to OVA in our behavioural paradigm, and whether mast cells were necessary for these effects, we sensitized WT and littermate control RMB mice and depleted mast cells with diphtheria toxin as described (Fig. 2g). Mast cell-depleted mice were then injected with increasing doses of recombinant GDF15 immediately prior to each day of OVA preference testing. We found that GDF15 rescued allergen aversion in mast cell-depleted mice, consistent with GDF15 being downstream of mast cell activation (Fig. 4d). This aversive response was dose dependent, leading to a partial aversive response at 0.01 mg/kg and one comparable with sensitized BALB/c mice at 0.1 mg/kg. The doses necessary to drive OVA aversion were higher than that induced by intragastric OVA challenge (Fig. 4 a, b). Thus, we sought to directly test whether GDF15 was necessary for the aversive response to OVA using anti-GDF15 neutralizing antibody^60^. We sensitized WT mice with OVA/alum and treated them 5 h prior to each day of OVA preference testing with isotype control or anti-GDF15 antibody (Extended Data Fig. 7a). We found GDF15 neutralization had no effect on the first day of two-bottle preference testing in sensitized mice (data not shown). On day 2 of the preference test, however, blocking GDF15 led to increased consumption of OVA in allergic sensitized mice relative to isotype control treatment (Fig. 4e, f). The increase in OVA preference could not be explained by differences in antibody titters, as the levels of total and OVA-specific IgE as well as OVA-specific IgG1 were similarly induced in isotype and anti-GDF15 treated groups (Extended Data Fig. 7b). Neutralization of GDF15 decreased preference to OVA in sensitized mice within 60 min and lasted throughout the 3 hours of the second trial (Extended Data Fig. 7c). GDF15 appears to be necessary for allergen aversion, yet it is incapable of driving aversion alone at concentrations comparable to the allergen challenge, suggesting that other signals may synergize with GDF15 to promote aversion. Leukotrienes are potential mediators that could explain this effect. Furthermore, our finding that GDF15 – but not IgE, mast cells, or leukotrienes – is dispensable for the first day of aversion suggests that leukotrienes may act through an acute pathway that GDF15 later sustains. Thus, we here describe an unexpected mast cell epithelial circuit that generates allergen aversion through a leukotriene- and GDF15-dependent mechanism. Together, these findings demonstrate that immune sensing of allergens leads to the generation of aversive behaviour. We suggest that avoidance behaviour is a defence strategy aimed at minimizing harmful effects of exposure to harmful substances, including allergens. Our findings suggest that detection of allergens by IgE antibodies expands sensory capacity of the nervous system and provides a mechanism to evaluate the quality of the food and other environmental factors. This conclusion adds to a growing body of evidence indicating that the immune detection of noxious stimuli is an important source of sensory information that drives corresponding behaviours^61^. It also adds to the growing evidence of by-directional functional interactions between the immune and nervous systems^62,63^.

## METHODS

### Animals

All animal care and experimentation were approved by the Institutional Animal Care and Use Committee of Yale University School of Medicine and consistent with the National Institutes of Health, USA, guidelines. Mouse lines were interbred in our facilities to obtain the final strains described in the text. Genotyping was performed according to the protocols established for the respective strains by The Jackson Laboratories or published by the donating investigators. Mice were maintained at the Yale University animal facilities in temperature- (22°C) and humidity-controlled rooms, in a 12 h light/dark cycle with free access to standard chow diet (Teklad 2018S, Envigo) and water.

Female mice at 6-10 weeks of age were used for all experiments. BALB/cJ (000651), C57BL/6J (000664), C57BL/6 FcεRI KO (B6.129S2(Cg)-*Fcer1a*^*tm1Knt*^/J, 010512), DdblGATA1 (C.129S1(B6)-Gata1^tm6Sho^/J, 005653)^64^, BALB/c Il4ra KO (BALB/c-*Il4ra*^*tm1Sz*^/J, 003514)^65^, and C57BL/6 substance P KO (B6.Cg-*Tac1*^*tm1Bbm*^/J, 004103)^44^ mice were purchased from The Jackson Laboratories and maintained in our facilities. BALB/c IgE KO were generously provided by H. C. Oettgen (Harvard University), RMB (B6. Ms4a2^tm1Mal^) were generously provided by P. Launay (Université Paris Diderot), and C57BL/6 *Trpm5-/-* mice were provided by W. Garret (Harvard University). RMB, FcεRI KO, IgE KO, or Trpm5-/- mice were backcrossed more than eight times onto BALB/cJ or C57BL/6J for this study. We used littermate controls in all experiments.

Chimeric mice were generated following a standard protocol^66^. C57BL/6J WT or FcεRI KO mice were used as donors in bone marrow (BM) transplant experiments. C57BL/6J WT recipient mice underwent a lethal total-body irradiation with 2 doses of 500 rad (Gammacell 40^137^Cs g-irradiation source), with an interval of 3 hours between the first and second irradiations. Fresh, unseparated BM cells (10 × 10^6^ per mouse) were injected into the tail vein of the irradiated recipient mice 4 hours after lethal irradiation. Chimerism efficiency was checked by flow cytometry 8 weeks post-irradiation and transplant using peripheral blood, and reconstituted mice were used 2 months after BM transplantation.

For mast cell depletion, sensitized RMB mice were injected i.p. with 0.05 mg/kg of diphtheria toxin (Sigma Aldrich D0564) three times every other day as previously described^35^, starting on day 28 after the first allergic sensitization. Because this protocol was efficient at depleting mast cells even in heterozygous RMB mice, we used both mutants and heterozygotes as mast cell-depleted models.

For drug trials, famotidine (Sigma-Aldrich F6889) and loratadine (Sigma-Aldrich, L9664) were both i.p. injected to a final concentration of 10 mg/kg for two days prior to the preference test. p-Chlorophenylalanine (Tocris, 0938) was injected i.p. for 5 consecutive days prior to the preference test to a final concentration of 100 mg/kg. Zileuton (Tocris, 3308) was used at 50 mg/kg in 0.6% methylcellulose by gavage, 1 h prior to each day of preference testing. Ondansetron hydrochloride (Tocris, 2891) was injected i.p. at 1 mg/kg 1 h before preference test. Aprepitant (Sigma-Aldrich SML2215) was i.p. injected 1 h prior to preference test at 5 mg/kg. Indomethacin (Sigma-Aldrich, I7378) was orally administered in the drinking water for 5 consecutive days prior to the preference test at a final concentration of 0.02 mg/mL. The CGRP selective antagonist, BIBN 4096 (Tocris, 4561), was i.p. injected at 0.4 mg/kg 50 min before preference test. All drugs were solubilized following the manufacturer’s instructions. All control groups received the appropriate vehicle solutions

For antibody treatment experiments, animals were injected i.v. through the retro-orbital sinus with control antibody against keyhole limpet hemocyanin (clone LTF-2) and GDF15 blocking antibody (Patent ID: WO2014100689A1, kind gift of Dr. Hui Tian)^67^ both at 10 mg/kg in 0.1 mL of PBS 6 h before preference test. Purified mouse GDF15 (Patent ID: WO2012138919A2) was injected i.p. 1 h prior to the preference test at concentrations ranging from 0.001 to 0.1 mg/kg.

### Allergic sensitization and challenges

Animals were sensitized subcutaneously on days 0 and 7 with 0.25 mg/kg endotoxin-free ovalbumin (OVA, BioVendor 321001) adsorbed in 50 mg/kg aluminium hydroxide gel (alum, Invivogen vac-alu-250) and diluted to a final volume of 0.2 mL in phosphate buffered saline (PBS) pH=7.4. Controls received all the above except for OVA (referred as PBS/alum).

For oral sensitization experiments, mice were given oral gavages on days 0 and 7 with 5 mg of grade III OVA with 0.5 mg/kg of cholera toxin (List Biologicals 100B, lot nos. 10165A1 and 10165A2) in a final volume of 0.25 mL 0.2 M sodium bicarbonate buffer. Both sensitization and challenge were performed around ZT4.

For allergen challenges, all groups received 5 oral gavages with 40 mg grade III OVA (Sigma-Aldrich A5378) in 0.25 mL of regular drinking water on days 14, 18, 21, 24, and 28, unless otherwise stated. For Figures 3 and 4 and extended data 5 and 6, all groups received 5 oral gavages with 50 mg grade III OVA (as above) in 0.2 mL of PBS on days 14, 16, 18, 20, 22.

### Preference test

Drinking behaviour was determined using the two-bottle preference test in custom-built lickometer cages. Two days after the last sensitization, on day 9, mice were individually placed into the lickometer cages and transferred to the temperature-controlled test room. During the 5 days of adaptation, mice were provided continuous exposure to two regular water bottles. Baseline measurements started on the last day of adaptation, day 14. On day 15, each mouse was given one bottle of water and one bottle containing 1% OVA (grade II, Sigma A5253), unless otherwise stated, 30 min before the lights turned off. The number of licks in each bottle was recorded periodically (1 or 5 min, as indicated) overnight. On day 16 at ZT2, OVA bottles were switched back to regular water bottles and the baseline preference was measured until the next day. On day 17, mice were provided with two new bottles 30 min before the lights turned off: one containing water and another containing a fresh solution of 1% OVA. The bottles’ positions were switched to account for potential place preference. Preference results are expressed as cumulative licks over time or as percentage OVA preference, calculated using the area under the curve of cumulative licks from the OVA bottle divided by the total cumulative licks (OVA + water). Total solution intake was also measured by calculating the difference in solution weight before and after the preference test. Mice had *ad libitum* access to chow diet.

Aversion specificity was tested using bovine serum albumin (BSA, Fisher BP1600-1). We used denatonium benzoate (DB, Sigma D5765) as the bitter compound for positive control of aversion. For one-bottle tests, mice were offered 1% OVA solution as their only drinking source for seven consecutive days. OVA solutions were prepared fresh each day. The difference in bottle weight between days was used to determine the drinking preference of control and sensitized animals.

### Brain tissue preparation and immunohistochemistry

Brains were collected 90 min after the first oral challenge with OVA. To minimize gavage-induced stress, we administered sham gavages with regular drinking water for 3 days before the final allergen challenge. Mice were deeply anesthetized with isoflurane (Covetrus) and were transcardially perfused with PBS followed by freshly prepared 4% paraformaldehyde (PFA) in PBS. Dissected brains were kept in 4% PFA at 4°C for 48 h, washed 3 times in PBS and transferred to a 30% sucrose in PBS solution for 2 days and then sliced into 40 um-thick coronal sections (AP, DMV, NTS, and PBN) and 100 um-thick sections (LH and CeA) using a Leica CM3050 S cryostat (Leica Biosystems). Briefly, the sections were permeabilized with PBS/0.3% Triton-x-100 for 30 minutes at room temperature, then blocked in PBS, 0.3% Triton-X-100, and 10% normal donkey serum in 0.3M glycine for 1 h at room temperature. Blocking was followed by incubation with rabbit monoclonal anti-cFos primary antibody (1:1000 dilution, Cell Signaling 2250S) or rabbit polyclonal anti-Fluoro-Gold primary antibody (1:1000 Fluorochrome) in the same blocking solution overnight for 16 h and then with Alexa Fluor 594-conjugated donkey anti-rabbit IgG secondary fluorescent antibody (1:500 dilution, Invitrogen A21207) for 2 h at room temperature. After being washed with the permeabilization solution again, the sections were mounted on slides and visualized by using a fluorescent All-In-One Keyence microscope (model BZ-X710, Keyence Corporation of America). Images were taken using the 4x objective. Brains regions were defined based on the Allen Mouse Brain Atlas reference atlas (mouse.brain-map.org/) and processed using the open-source Fiji-ImageJ software^68^. Fos-positive cells were manually quantified by a blinded investigator throughout the entire procedure.

### Antibody quantification

Serum levels of total IgE and OVA-specific IgE antibodies were determined by sandwich ELISA. For total IgE, ELISA-grade plates (490012-252, VWR) were coated overnight at 4°C with 2 µg/mL of anti-mouse IgE (553413, clone R35-72, BD Pharmingen) in 0.1 M sodium carbonate buffer pH 9.5. Plates were blocked with 1% BSA (Fisher BP1600-1) for 2 h at room temperature. Serum from sensitized and control mice was diluted up to 1:100 and incubated for 2 h at room temperature. Purified mouse IgE (BD Biosciences 557079 for BALB/c and 557080 for C57BL/6) was used as standard at the highest concentration of 100 ng/mL followed by two-fold dilutions to create a standard curve. Afterwards, 500 ng/mL of biotin-conjugated anti-IgE detection antibodies (553419, clone R35-118, BD Biosciences) were incubated for 1 h at room temperature followed with another incubation of diluted HRP-conjugated streptavidin (554066, BD Biosciences) for 30 min at room temperature. Then plates were incubated in the dark at room temperature with TMB substrate reagent (555214, BD Biosciences) and the colour was checked every 3 min. Plates were read at 450 nm immediately after adding stop solution (3M H_2_SO_4_). Between each step, plates were washed five to seven times with 0.05% Tween-20 in PBS. Serum concentrations of OVA-specific IgE were assayed by the same ELISA method. Purified mouse anti-OVA IgE (MCA2259, clone 2C6, BioRad) was used as standards with the highest concentration at 100 ng/mL. Capture antibodies were the same as for total IgE assay and 8 mg/mL of biotinylated OVA (OVA1-BN-1, Nanocs) was used for detection. OVA-specific IgG1 antibodies in the serum were assayed by direct ELISA. Plates were coated overnight at 4°C with 20 µg/mL of grade V OVA (A5503, Sigma-Aldrich) in coating buffer (0.1 M sodium carbonate pH 9.5). After the blocking step, serum samples were diluted up to 1:10,000 and incubated for 1 hr at room temperature. Purified mouse BALB/c IgG1 (557273, clone MOPC-31C, BD Biosciences) was used as standard with the highest concentration at 100 ng/mL. Biotin-conjugated mouse IgG1 (553441, clone A85-1, BD Biosciences) was used for detection at 100 ng/mL. The HRP-conjugated streptavidin, substrate reagents, blocking and washing solutions used for IgG1 ELISA were the same as described above for IgE ELISAs.

### Oral and systemic anaphylaxis

To determine the occurrence of active systemic anaphylaxis, sensitized mice were challenged intravenously (i.v.) through the retro-orbital sinus with 5 mg/kg of grade V OVA (A5503, Sigma-Aldrich) 14 days after the first sensitization at ZT4. For oral anaphylaxis, mice were administered with 40 mg of grade III OVA (A5378, Sigma-Aldrich) intragastric. Rectal temperature was measured every 30 min for 4 h after challenge using a probe (Thermalert TH-5).

### Complete subdiaphragmatic vagotomy

Mice were sensitized with OVA and alum on days 0 and 7 as described above. Mice were placed on a liquid diet four to five days prior to promote survival and recovery. Mice were i.p. treated with bupivacaine (2 mg/kg), buprenorphine (1 mg/kg), and meloxicam (5 mg/kg). Mice were anesthetized with 4% isoflurane for induction and then transferred under the microscope and maintained at 1-1.5% isoflurane during the surgery. An abdominal midline incision was made through the skin and the intraperitoneal wall. The stomach was exteriorized to expose the oesophagus. Two blunted and bent 19G needles were gently placed at the proximal and distal end of the oesophagus to isolate it from the remaining tissue. The vagus nerves could be observed running dorsal and ventral to the oesophagus. The nerves were incised at the most proximal end of the oesophagus using curved fine forceps. Control sham-operated mice received the same surgical procedure except for the incision of the nerves. The blunted needles were removed, and the intraperitoneal cavity and the skin layer were sutured. Post-surgery animals were given a liquid diet for ten days with moistened chow also provided five days post-surgery. Animals were then placed on a regular chow diet and used for experimental purposes two weeks post-surgery. Histological verification of vagotomy was confirmed using an i.p injection of 0.1% Fluoro-Gold (Fluorochrome), followed by examination of its presence in the dorsal motor nucleus of the vagus (DMX) two weeks after injection.

### Gastrointestinal motility

Gastrointestinal (GI) transit time was assessed immediately following the fourth oral challenge with intragastric OVA, on day 24 after allergic sensitization at ZT3. Mice were gavaged with a 0.25 mL solution with 6% carmine red (C1022, Sigma-Aldrich), 0.5% methylcellulose (M0512, Sigma-Aldrich), and 40 mg grade III OVA. After oral gavage and for the duration of the assay, mice were individually placed in clean cages containing regular bedding. Mice had free access to food and water and were monitored for the occurrence of diarrhoea. The GI transit time was measured as the time between oral gavage and the appearance of the first faecal pellet containing the red carmine dye. Mice were grouped together at the end of the assay.

### Epithelial cell and lamina propria isolation

Single-cell suspensions of small intestinal epithelium and lamina propria were prepared as described^69,70^. Briefly, the small intestine was isolated and opened longitudinally. Its contents were then rinsed in PBS following removal of Peyer’s patches. The tissue was then cut into 2-3-cm segments and incubated in RPMI media (ThermoFisher) containing 5 mM EDTA, 0.145 mg/mL DTT, and 3% FBS at 37°C with 5% CO_2_ for 20 min with agitation. Pieces of intestine were washed in RPMI containing 2 mM EDTA to separate the epithelial fraction. This fraction was then subjected to a 30% Percoll density gradient by centrifugation (GE17-0891-01, Sigma-Aldrich). Lamina propria digestion was performed using 0.1 mg/mL Liberase TL (Roche #540102001) and 0.5 mg/mL DNAse (DN25, Sigma-Aldrich) in RPMI for 30 min at 37°C with 5% CO_2_. Digested tissue was sequentially strained through 70 µM and 40 µM strainers, washed in RPMI containing 3% FBS, and cells were then stained for further analysis.

### Flow cytometry

Single-cell suspensions were treated with anti-CD16/32 (Fc block) (14-9161-73, ThermoFisher) and stained with the live/dead marker ethidium monoazide bromide (ThermoFisher #E1374) in 2% FBS in PBS. The following antibodies were used at a concentration of 1 µg/mL except where otherwise indicated: APC-Cy7-CD117 (clone 2B8; Biolegend #105826), PE-FceRI (clone MAR-1; eBioscience 12-5898-82), eFluor450-CD45 (clone 30-F11; eBioscience #48-0451-82), APC-CD11b (clone M1/70; ThermoFisher #17-0112-82), Alexa700-CD19 (clone 6D5; Biolegend #B189284), PE/Cy7-CD3e (clone 145-SC11, eBioscience #25-0031-82), APC-MHC II (clone M5/114.15.2, eBioscience #17-5321-82), and PE-Siglec F (clone E50-2440, BD Pharmingen #5521126). Cells were fixed with 1.6% paraformaldehyde (Electron Microscopy Sciences, #15710). Flow cytometry was performed using a BD LSRII analyser equipped with the following lasers: 355 nm (UV), 405 nm (violet), 488 nm (blue), and 633 nm (red). Data was analysed using FlowJo software. Gates were drawn according to fluorescence minus one (FMO) controls.

### RNA Scope

For RNAscope experiments, FISH was performed on FFPE tissues as previously described (Wang, J Mol Diagn, 2012). Briefly, mice were sensitized as above and challenged i.g. six times every other day with 50 mg of grade III OVA. One hour after the sixth gavage, mice were euthanized by CO2 asphyxiation and small intestine and colon harvested. Tissues were flushed with PBS, followed by 4% PFA, opened longitudinally and fixed overnight at room temperature. The following day, the small intestine was divided into thirds, and all tissues were rolled using the Swiss roll technique and subsequently embedded in paraffin. RNAscope was performed using RNAscope Multiplex Flourescent v2 kit (ACD 323110) using probes specific for Gdf15 (ACD 318521), EPCAM (ACD 418151-C2) and Fcer1a (ACD 511331-C3) following manufacturer’s instructions for FFPE tissues with fluorescent detection via Opal 620, Opal 520, and Opal 570 respectively (Akoya Biosciences FP1495001KT, FP1487001KT, SKU FP1488001KT). Slides were counterstained with DAPI. Quality of tissue RNA was confirmed, and background threshold established using positive and negative control probes (ACD 320881, ACD320871). Images were acquired on a Leica STP 6000 Fluorescent Microscope using tile-scans of entire Swiss roll sections at 20x resolution. FceR1a, EPCAM, and GDF15 colocalization analysis was performed in Qupath (<u>QuPath | Quantitative Pathology & Bioimage Analysis</u>) based on fluorescent threshold of cells detected by DAPI positivity.

### GDF15 quantification

Unless otherwise noted, serum for all GDF15 measurements were taken from mice 1 h after 50 mg i.g. intragastric OVA challenge. GDF15 levels in the serum were measured by ELISA (R&D, MGD150).

### RNA extraction and quantification

For tissue RNA extraction, tissues were harvested into RNA STAT-60 RNA isolation reagent (Amsbio) and disrupted by bead homogenization (Omni, INC). RNA was extracted using the Direct-zol RNA Mini Kit (Zymo, R2051). cDNA synthesis was performed using MMLV reverse transcriptase (Clontech) with oligo(dT) primers. qRT PCR was performed on CFX384 Real-Time System (Bio-Rad) using PerfeCTa SYBR Green Supermix (Quanta Biosciences) and transcripts were normalized to *Rpl13a*.

### Statistics

Statistical analyses were performed in GraphPad Prism software. Data were analysed with Mann-Whitney *U* test (two experimental groups) or Dunnett’s multiple comparison test. Statistical significance is defined as *p<0.05, **p<0.01, ***p<0.001. Nonparametric statistical analyses were used throughout the manuscript, unless stated otherwise. All data are mean ± s.e.m.

## Acknowledgements

We thank all Medzhitov and Dietrich Lab members for discussions. C. Annicelli, S. Cronin, and J. Bober for their assistance with animal care and husbandry. We thank W. Khoury-Hanold for backcrossing the *Trpm5*-/-into BALB/c background. We thank M. Zimmer for discussions and for his technical assistance with behavioural assays. We thank D. Mucida for fruitful discussions and for providing us with the RMB F1 mice. We also thank T. Carvalho for constructive suggestions. Funding: The R.M. laboratory is supported by the Howard Hughes Medical Institute, Food Allergy Science Initiative, Blavatnik Family Foundation, and a grant form NIH (AI144152-01). R.M. is an investigator of the Howard Hughes Medical Institute. N.D.B is supported by the National Institute of Allergy and Infectious Diseases of the National Institutes of Health under Award Number F30AI174787.

## Author contributions

E.B.F. and R.M. conceived and designed the study. M.D. and A.W. contributed ideas and expertise that impacted the goals of this study. E.B.F., N.D.B., and J.C. performed the preference tests and inflammation experiments and generated the BALB/c RMB, and C57BL/6 IgE KO mice. E.B.F. and M.D. built the initial lickometer cages and established the preference tests. F.C. and B.G. performed the brain collections, cFos immunostaining, and quantifications. M.G. performed the vagotomy surgeries and preference tests with vagotomised mice. C.Z. generated the FcεRI chimeras. G.G. and P. L. developed the RMB mice. E.B.F., N.D.B., J.C., B.G., M.G., and M.D. analysed data. E.B.F., N.D.B, and R.M. wrote the manuscript with input from J.C, A.W. and M.D. All authors revised and edited the manuscript and figures.

## Extended Data Figures

**Extended Data Fig. 1.**
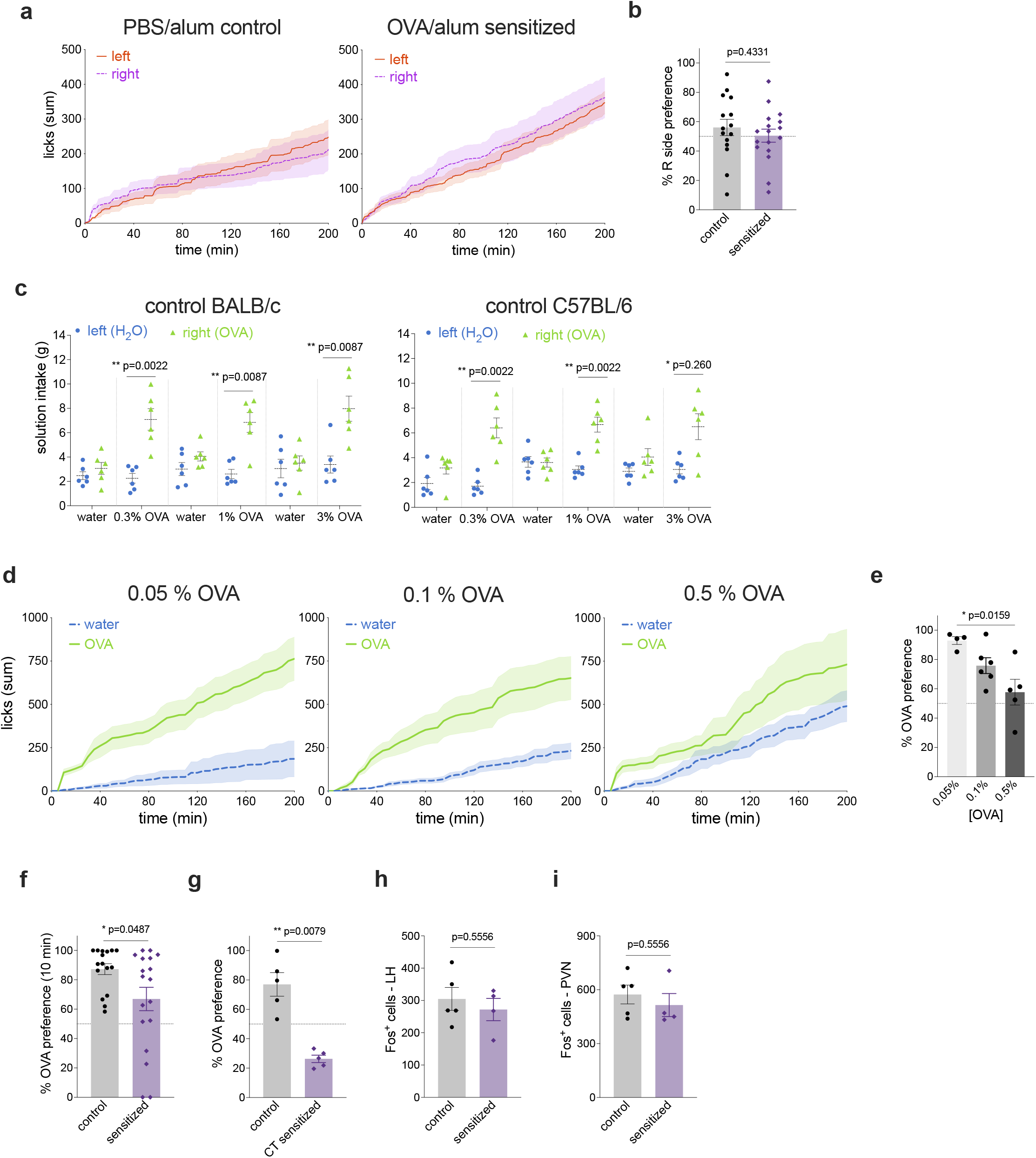
Characterization of immunological aversion. **a**, Cumulative licks of water solution from right- or left-positioned bottles from control (left) and OVA/alum sensitized mice (right). **b**, Percentage preference of control and OVA/alum sensitized mice to the right side water bottle. **c**, Solution intake of water and varying concentrations of OVA in control BALB/c (left) and C57BL/6 (right) on day 1 of the two-bottle preference test. **d**, Cumulative licks to different concentrations of OVA from mice sensitized with OVA/alum. **e**, Preference to the bottle containing OVA in OVA/alum sensitized mice. **f**, Preference to 1% OVA solution by control and OVA/cholera toxin-sensitized mice. **h - i**, Total number of c-fos+ cells in the lateral hypothalamus (LH) and paraventricular nucleus of the hypothamlamus (PVN) of OVA/alum sensitized BALB/c mice.

**Extended Data Fig. 2.**
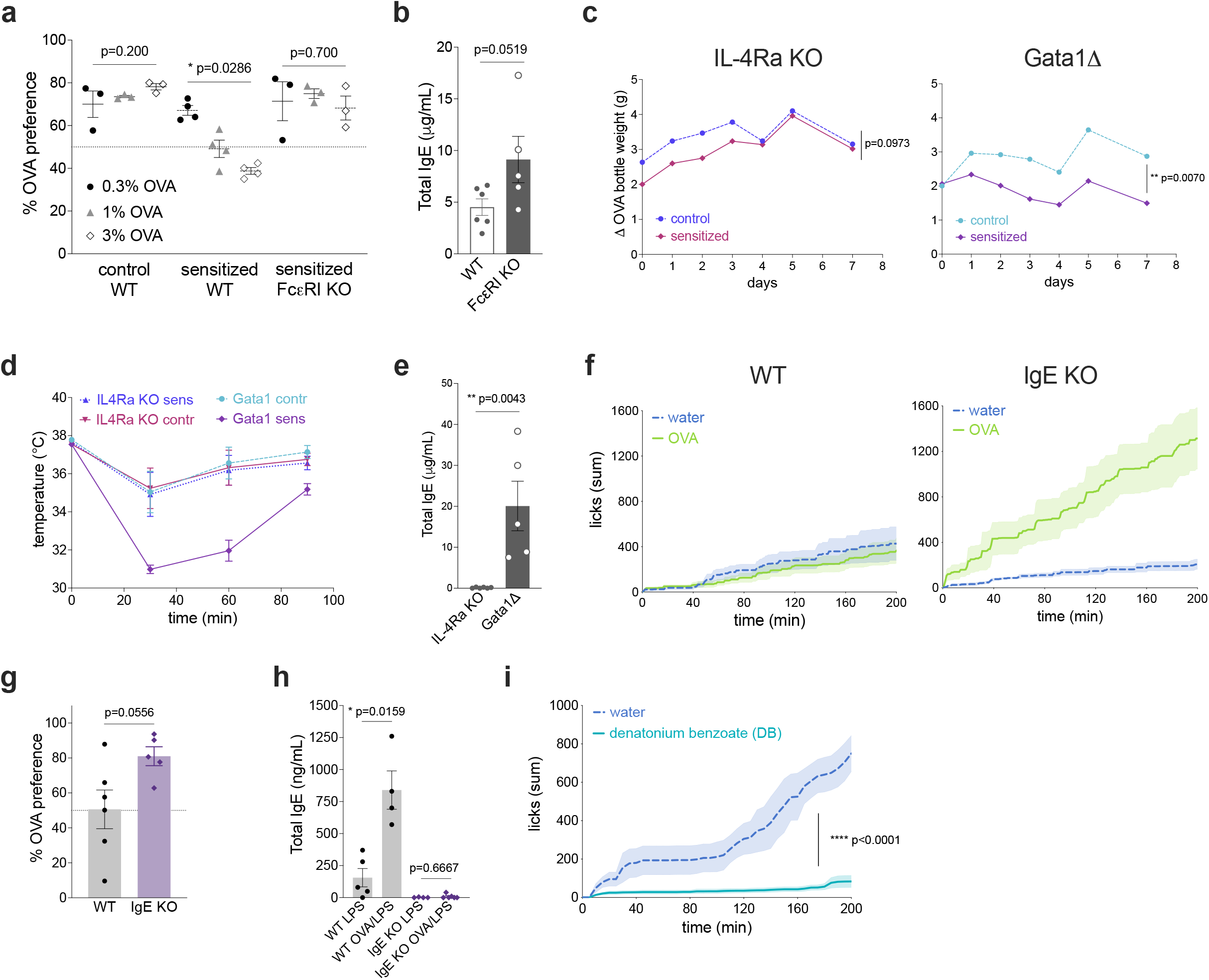
Role of IgE signaling in allergic and non-allergic aversion. **a**, Drinking preference of C57BL/6 WT and FcεRI KO to OVA by solution intake of different concentrations of OVA. **b**, Total serum IgE in WT and FcεRI KO mice sensitized with OVA/alum and orally challenged five times with OVA. **c**, Solution intake of OVA was measured for one week in BALB/c IL-4Ra KO (left) or Gata1Δ (right) mice orally sensitized with OVA/CT. **d**, Rectal temperature after systemic OVA challenge in OVA/CT orally sensitizatized mice. **e**, Total levels of serum IgE after OVA/CT sensitization and five oral challenges with OVA. **f**, Two-bottle preference test in C57BL/6 WT (left) or IgE KO (right) mice sensitized with OVA and LPS. **g**-**h**, Preference to OVA (**g**) and total serum IgE (**h**) in OVA/LPS sensitized WT and IgE KO mice. **i**, Cumulative licks during the two-bottle preference test with water and a solution with the bitter compound denatonium benzoate (DB) in C57BL/6 IgE KO mice sensitized with OVA/ alum.

**Extended Data Fig. 3.**
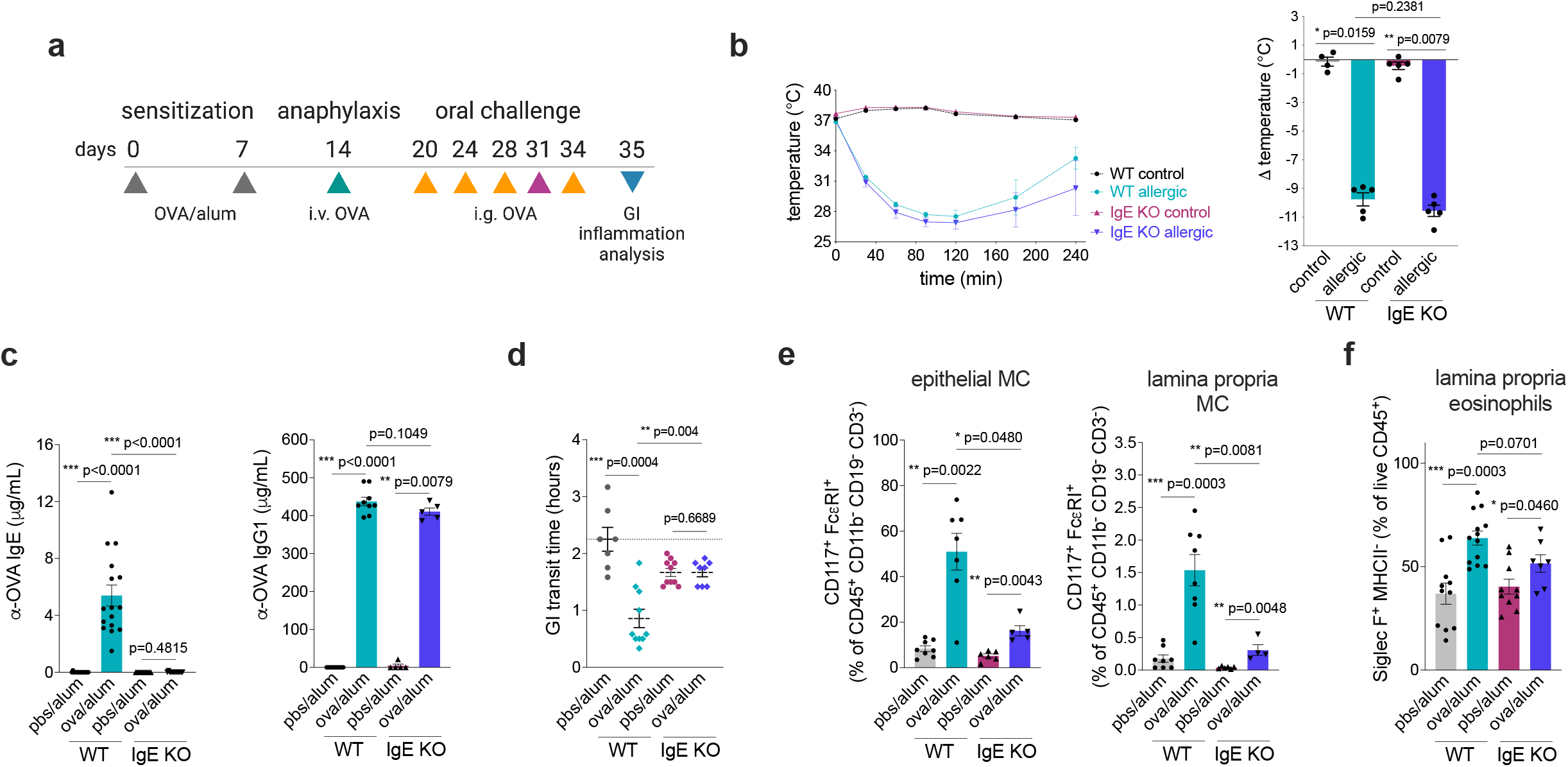
IgE-dependent and independent roles in gut allergic inflammation. **a**, Experimental protocol for the development of gut allergic inflammation and systemic anaphylaxis. **b**, Rectal temperature over time (left) and maximum temperature variation (right) after systemic OVA challenge on day 14 in OVA/alum sensitized BALB/c WT and littermate IgE KO mice. **c**, Serum specific IgE (left) and IgG1 (right) antibodies in sensitized BALB/c WT and IgE KO mice after five oral challenges with OVA, on day 35. **d**, Gastrointestinal transit time in BALB/c WT and IgE KO mice determined with red carmine assay on day 31 upon oral allergen challenge. **e**-**f**, Accumulation of mast cells and eosinophils in the small intestines of sensitized BALB/c WT and IgE KO mice on day 35.

**Extended Data Fig. 4.**
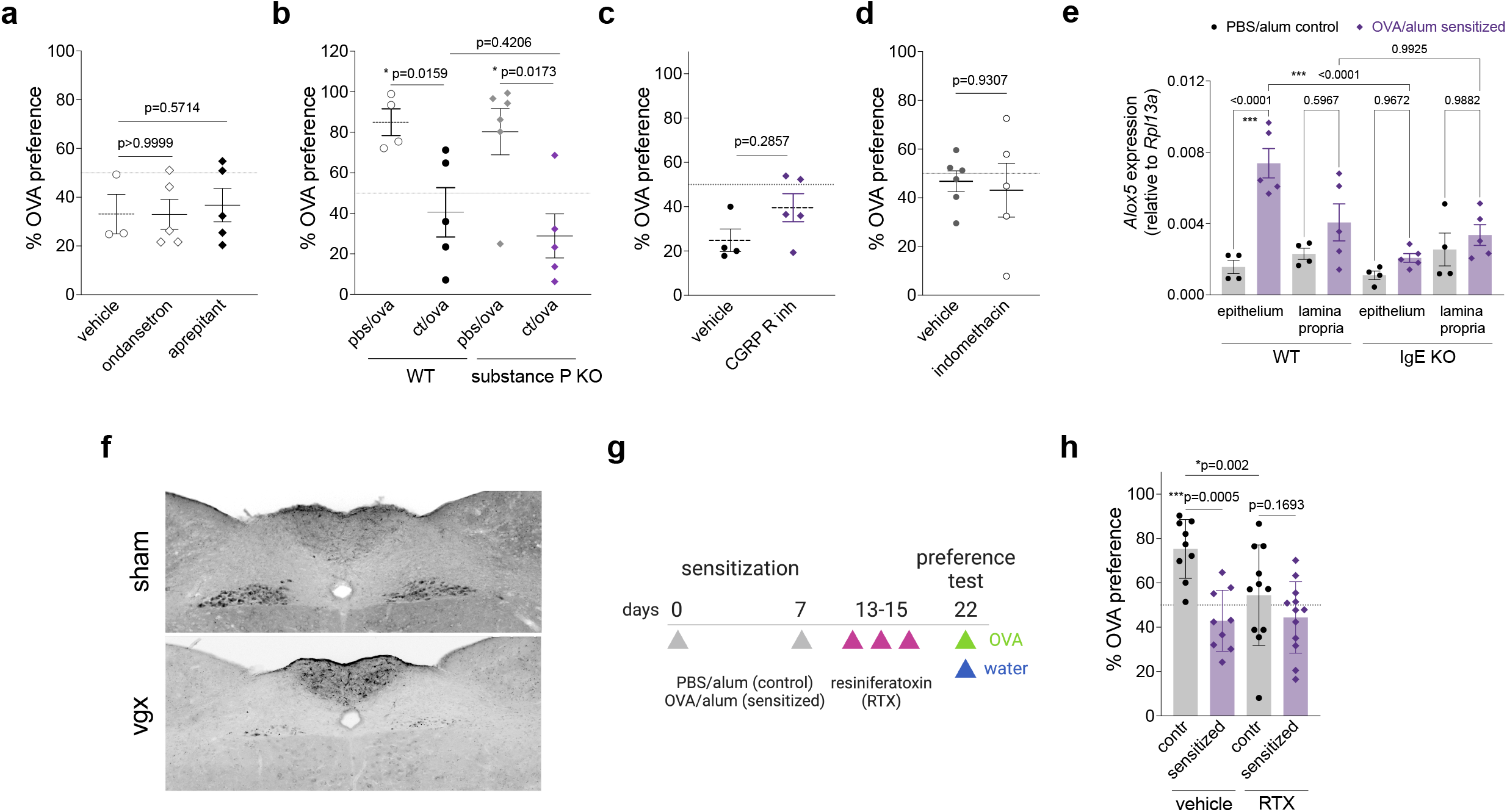
Allergen-induced avoidance behavior is independent on substance P, CGRP, and prostaglandins. **a, c**, Drinking preference to OVA solution by OVA/alum sensitized BALB/c mice that were i.p. injected with the serotonin receptor 3 antagonist (ondansetron), substance P receptor NK1 antagonist (aprepitant), CGRP receptor inhibitor (BIBN4096) or vehicle solution prior to preference test. **b**, Preference to OVA in C57BL/6 WT or substance P KO mice orally sensitized with OVA and cholera toxin. **d**, OVA preference in OVA/alum sensitized BALB/c mice administered with the cyclooxygenase inhibitor indomethacin or vehicle solution in the drinking water for 5 days prior to the preference test. **e**, *Alox5* expression levels in the epithelial (Ep) and lamina propria (LP) compartments of the small intestine from BALB/c WT or IgE KO mice sensitized with PBS/alum (control) or OVA/alum (sensitized). n=4-5 per group. **f**, Staining of the dorsal motor nucleus of the vagus after fluorogold injection into the small intestine of WT mice to confirm efficiency of subdiaphragmatic vagotomy. Representative **g**, For TRPV1+ cell depletion, BALB/c mice were sensitized with OVA/alum, as indicated. On week 3, mice were i.p. injected with resiniferatoxin (RTX) every other day for 3 days and the two-bottle preference test was assessed one week after the last RTX injection. **h**, Preference to OVA solution in vehicle- or RTX-treated mice. **a** - **d, h**, Mean ± s.e.m., Mann-Whitney test. **e**, Mean ± s.e.m., One-Way ANOVA with multiple comparisons. **a** - **h**, Representative of two experiments.

**Extended Data Fig. 5.**
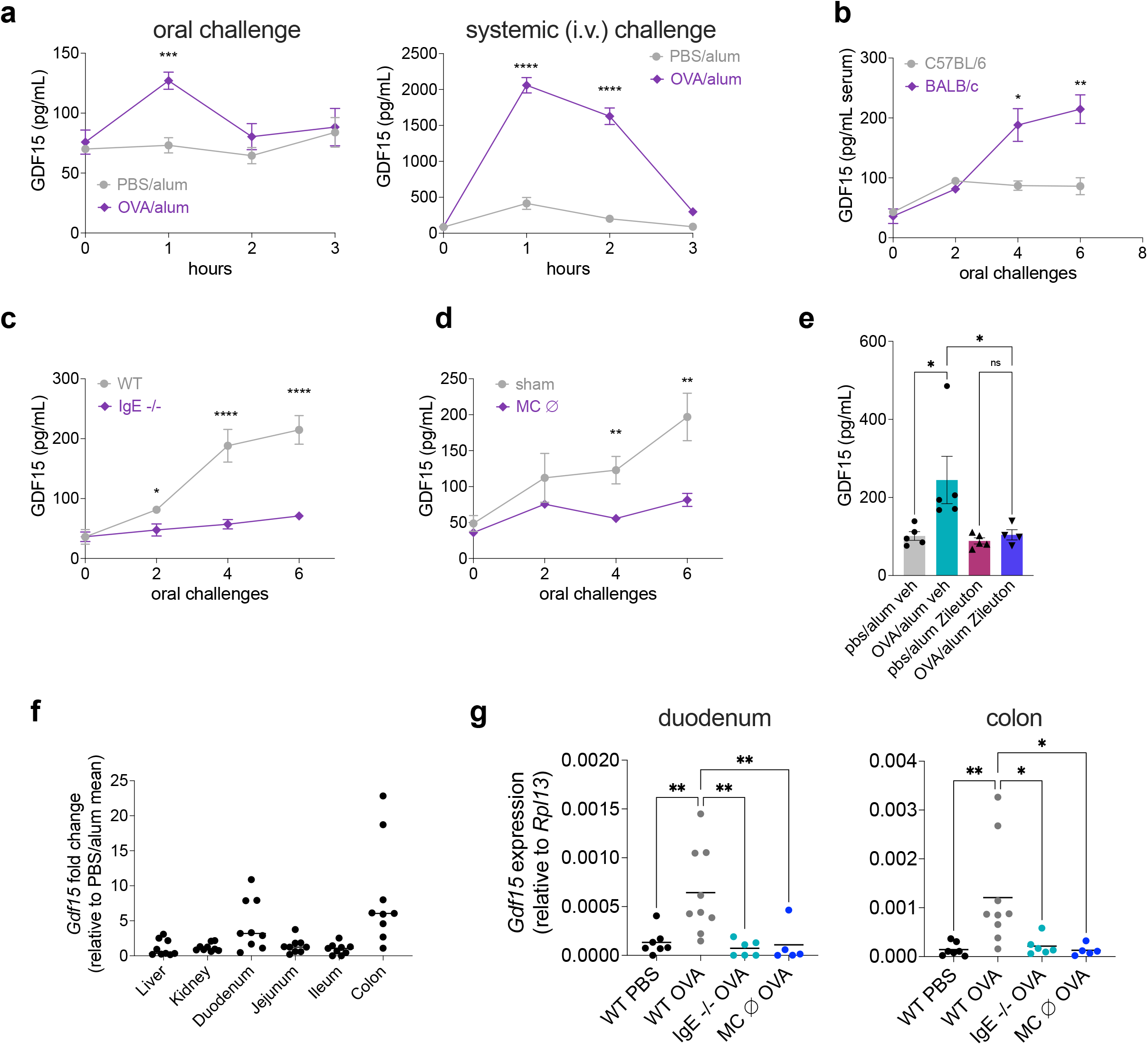
GDF15 is induced upon allergen exposure. **a**, GDF15 serum levels induced by oral (left) or intravenous (right) OVA administration in sensitized BALB/c mice. **b**-**d**, Serum GDF15 induction after oral allergen challenges in OVA/alum sensitized BALB/c and C57BL/6 (b), IgE KO (c), or mast cell-depleted RMB (d) mice. **e**, Induction of serum GDF15 in control and sensitized mice after the administration of the 5-lipoxygenase inhibitor zileuton during oral challenges with OVA. **f**, Fold change in *Gdf15* mRNA transcripts in the intestinal tissues of OVA/alum sensitized BALB/c mice relative to control groups. **g**, *Gdf15* mRNA transcripts in the duodenum (left) and colon (right) in sensitized mice.

**Extended Data Fig. 6.**
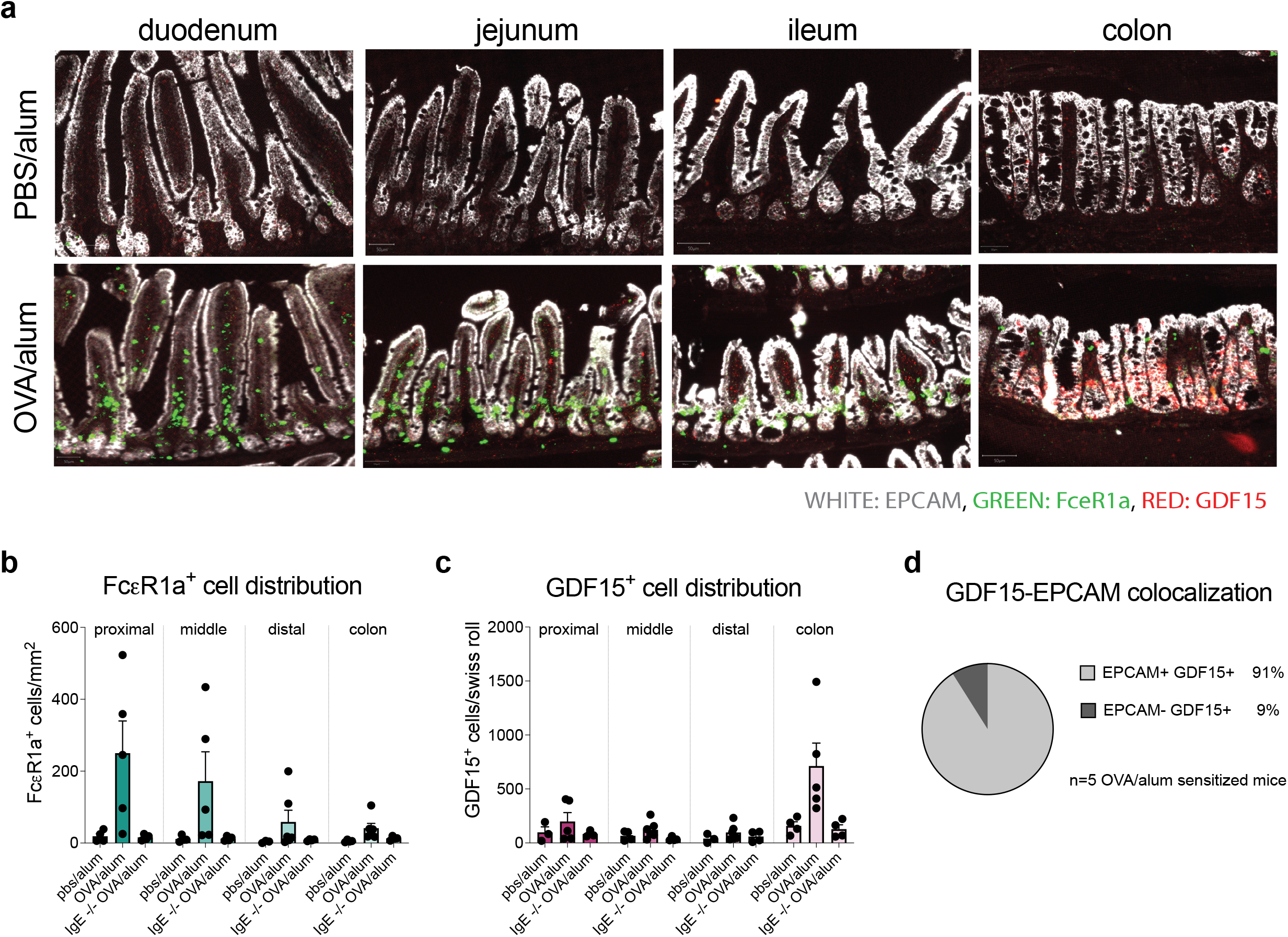
GDF15 originates from colonic epithelial cells. **a**, Fcεr1a (green), EPCAM (grey), and GDF15 (red) transcripts across intestinal tissues in OVA/alum BALB/c sensitized and control mice by RNAScope. Magnitude of images is 200x. **b-c**, Analysis of intestinal distribution of FcεR1 expressing cells (b) and GDF15 expressing cells (c) from control and allergic sensitized WT or IgE KO mice. Quantification was performed after RNAscope technique. **d**, Colocalization analysis of GDF15 expressing colonic cells and cells expressing the epithelial cell marker, EPCAM.

**Extended Data Fig. 7.**
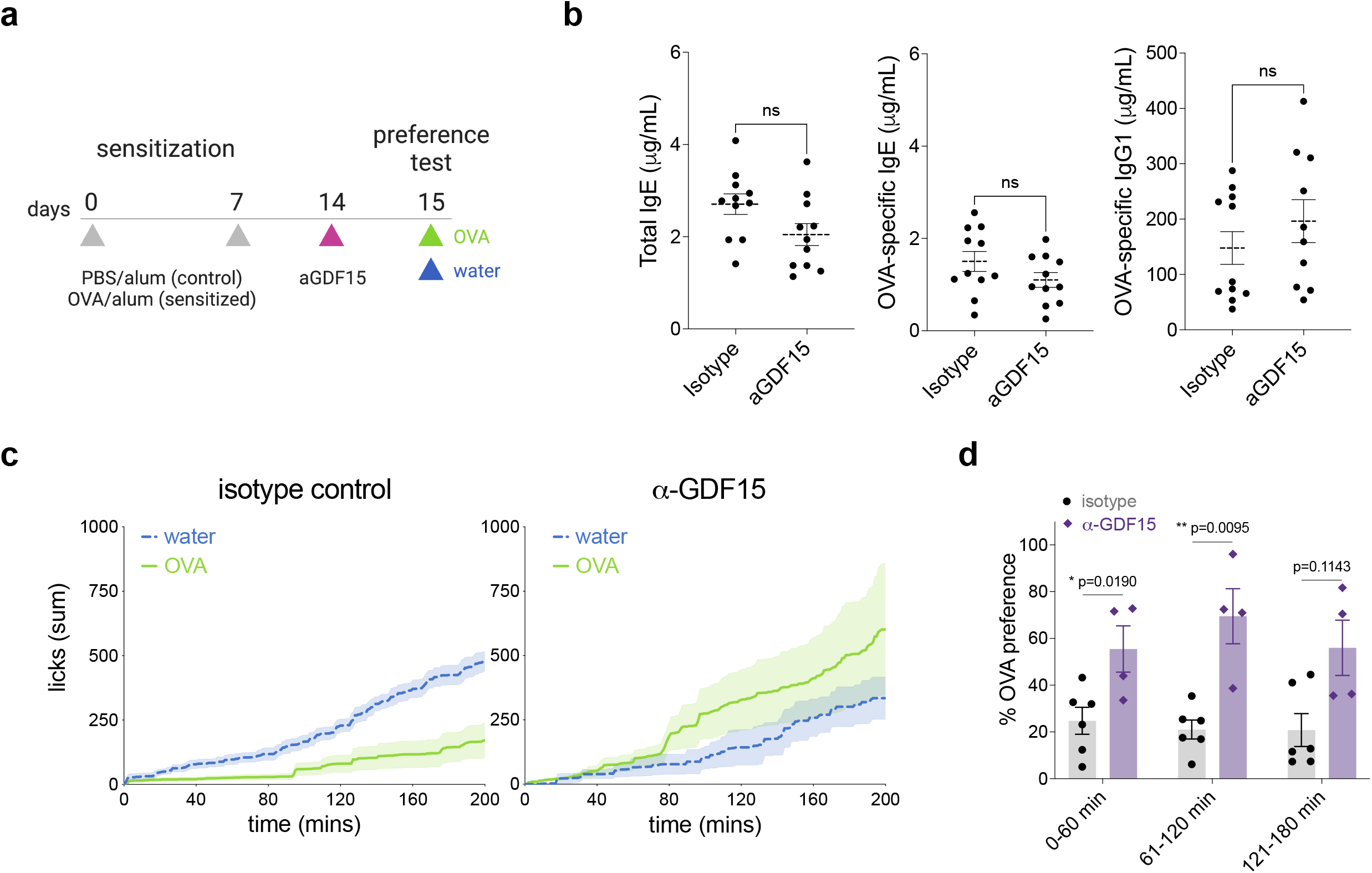
GDF15 blockade reduces allergen aversion. **a**, BALB/c mice were sensitized with OVA/ alum and treated with a mouse GDF15 blocking antibody or isotype control 5 hours prior to two bottle preference testing. **b**, Total IgE, OVA-specific IgE, and OVA-specific IgG1 antibodies after GDF15 antibody treatment. **c**, Cumulative licks during the two-bottle preference test of isotype control (left) or GDF15 blocking antibody-treated (right) mice. **d**, OVA preference of OVA/alum sensitized mice that received either GDF15 blocking antibody or isotype control 5 h prior to the behavioral assay.

## References

1. Hoffman, S. A., Shucard, D. W. & Harbeck, R. J. The immune system can affect learning: Chronic immune complex disease in a rat model. J. Neuroimmunol. 86, 163–170 (1998).

2. De Roode, J. C. & Lefèvre, T. Behavioral immunity in insects. Insects 3, 789–820 (2012).

3. Talbot, S., Foster, S. L. & Woolf, C. J. Neuroimmunity: Physiology and Pathology. Annu. Rev. Immunol. 34, 421–447 (2016).

4. Jackson, K. D., Howie, L. D. & Akinbami, L. J. Trends in Allergic Conditions Among Children : United States, 1997-2011. NCHS Data Brief 1–8 (2013).

5. Strachan, D. P. Hay fever, hygiene, and household size. BMJ 299, 1259–60 (1989).

6. Okada, H., Kuhn, C., Feillet, H. & Bach, J.-F. The ‘hygiene hypothesis’ for autoimmune and allergic diseases: an update. Clin. Exp. Immunol. 160, 1–9 (2010).

7. Schaub, B., Lauener, R. & von Mutius, E. The many faces of the hygiene hypothesis. J. Allergy Clin. Immunol. 117, 969–977 (2006).

8. Moneret-Vautrin, D. A., Morisset, M., Flabbee, J., Beaudouin, E. & Kanny, G. Epidemiology of life-threatening and lethal anaphylaxis: A review. Allergy Eur. J. Allergy Clin. Immunol. 60, 443–451 (2005).

9. Profet, M. The function of allergy: immunological defense against toxins. Q. Rev. Biol. 66, 23–62 (1991).

10. Stebbings, J. H. Jr. Immediate Hypersensitivity: A Defense against Arthropods? Perspect. Biol. Med. 17, 233–241 (1974).

11. Marichal, T. et al. A beneficial role for immunoglobulin E in host defense against honeybee venom. Immunity 39, 963–975 (2013).

12. Palm, N. W. et al. Bee venom phospholipase A2 induces a primary type 2 response that is dependent on the receptor ST2 and confers protective immunity. Immunity 39, 976–85 (2013).

13. Undem, B. & Taylor-Clark, T. Mechanisms underlying the neuronal-based symptoms of allergy. J. Allergy Clin. Immunol. 1–14 (2014).

14. Cara, D. C., Conde, A. A. & Vaz, N. M. Immunological induction of flavor aversion in mice. Braz. J. Med. Biol. Res. 27, 1331–1341 (1994).

15. Basso, A. S., De Sá-Rocha, L. C. & Palermo-Neto, J. Immune-induced flavor aversion in mice: Modification by neonatal capsaicin treatment. NeuroImmunoModulation 9, 88–94 (2001).

16. DJURIĆ, V. J. et al. Conditioned taste aversion in rats subjected to anaphylactic shock. Ann. N. Y. Acad. Sci. 496, 561–568 (1987).

17. Damak, S. et al. Trpm5 null mice respond to bitter, sweet, and umami compounds. Chem. Senses 31, 253–264 (2006).

18. Mirotti, L., Mucida, D., de Sá-Rocha, L. C., Costa-Pinto, F. A. & Russo, M. Food aversion: a critical balance between allergen-specific IgE levels and taste preference. Brain. Behav. Immun. 24, 370–5 (2010).

19. de Araujo, I. E. Circuit organization of sugar reinforcement. Physiol. Behav. 164, 473–477 (2016).

20. Campos, C. A., Bowen, A. J., Roman, C. W. & Palmiter, R. D. Encoding of danger by parabrachial CGRP neurons. Nature 555, 617–620 (2018).

21. Alhadeff, A. L. et al. A Neural Circuit for the Suppression of Pain by a Competing Need State. Cell 173, 140-152.e15 (2018).

22. Butler, R. K. et al. Activation of phenotypically-distinct neuronal subpopulations of the rat amygdala following exposure to predator odor. Neuroscience 175, 133–144 (2011).

23. Galli, S. J. & Tsai, M. IgE and mast cells in allergic disease. Nat. Med. 18, 693–704 (2012).

24. Shade, K. T. C. et al. Sialylation of immunoglobulin E is a determinant of allergic pathogenicity. Nature 582, 265–270 (2020).

25. Tepper, R. I. et al. IL-4 induces allergic-like inflammatory disease and alters T cell development in transgenic mice. Cell 62, 457–467 (1990).

26. Finkelman, F. D. et al. IL-4 is required to generate and sustain in vivo IgE responses. J. Immunol. 141, 2335–2341 (1988).

27. Chu, D. K. et al. Indigenous enteric eosinophils control DCs to initiate a primary Th2 immune response in vivo. J. Exp. Med. 211, 1657–1672 (2014).

28. Oettgen, H. C. & Burton, O. T. IgE receptor signaling in food allergy pathogenesis. Curr. Opin. Immunol. 36, 109–114 (2015).

29. Tordesillas, L., Berin, M. C. & Sampson, H. A. Immunology of Food Allergy. Immunity 47, 32–50 (2017).

30. Florsheim, E. B., Sullivan, Z. A., Khoury-Hanold, W. & Medzhitov, R. Food allergy as a biological food quality control system. Cell 1–15 (2021) doi:10.1016/j.cell.2020.12.007.

31. Oettgen, H. C. et al. Active anaphylaxis in IgE-deficient mice. Nature 370, 367–370 (1994).

32. Brandt, E. B. et al. Mast cells are required for experimental oral allergen-induced diarrhea. J. Clin. Invest. 112, 1666–1677 (2003).

33. Aguilera-Lizarraga, J. et al. Local immune response to food antigens drives meal-induced abdominal pain. Nature 590, 151–156 (2021).

34. Barbara, G. et al. Mast Cell-Dependent Excitation of Visceral-Nociceptive Sensory Neurons in Irritable Bowel Syndrome. Gastroenterology 132, 26–37 (2007).

35. Dahdah, A. et al. Mast cells aggravate sepsis by inhibiting peritoneal macrophage phagocytosis. J. Clin. Invest. 124, 4577–4589 (2014).

36. Veiga-Fernandes, H. & Mucida, D. Neuro-Immune Interactions at Barrier Surfaces. Cell 165, 801–811 (2016).

37. Wernersson, S. & Pejler, G. Mast cell secretory granules: armed for battle. Nat. Rev. Immunol. 14, 478–494 (2014).

38. Bellono, N. W. et al. Enterochromaffin Cells Are Gut Chemosensors that Couple to Sensory Neural Pathways. Cell 170, 185-198.e16 (2017).

39. Bhargava, K. P. & Dixit, K. S. Role of the chemoreceptor trigger zone in histamine-induced emesis. Br. J. Pharmacol. 34, 508–513 (1968).

40. Solinski, H. J. et al. Nppb Neurons Are Sensors of Mast Cell-Induced Itch. Cell Rep. 26, 3561-3573.e4 (2019).

41. R, S. Serotonin and GI clinical disorders. Neuropharmacology 55, (2008).

42. Shim, W.-S. et al. TRPV1 mediates histamine-induced itching via the activation of phospholipase A2 and 12-lipoxygenase. J. Neurosci. Off. J. Soc. Neurosci. 27, 2331–2337 (2007).

43. Wallrapp, A. et al. Calcitonin Gene-Related Peptide Negatively Regulates Alarmin-Driven Type 2 Innate Lymphoid Cell Responses. Immunity 51, 709-723.e6 (2019).

44. Cao, Y. Q. et al. Primary afferent tachykinins are required to experience moderate to intense pain. Nature 392, 390–394 (1998).

45. Reed, D. E. et al. Mast cell tryptase and proteinase-activated receptor 2 induce hyperexcitability of guinea-pig submucosal neurons. J. Physiol. 547, 531–542 (2003).

46. Hsieh, F. H., Lam, B. K., Penrose, J. F., Austen, K. F. & Boyce, J. A. T helper cell type 2 cytokines coordinately regulate immunoglobulin E-dependent cysteinyl leukotriene production by human cord blood-derived mast cells: profound induction of leukotriene C(4) synthase expression by interleukin 4. J. Exp. Med. 193, 123–133 (2001).

47. m, M., Kf, A. & Jp, A. The immediate phase of c-kit ligand stimulation of mouse bone marrow-derived mast cells elicits rapid leukotriene C4 generation through posttranslational activation of cytosolic phospholipase A2 and 5-lipoxygenase. J. Exp. Med. 182, (1995).

48. Lewis, R. A. et al. Prostaglandin D2 generation after activation of rat and human mast cells with anti-IgE. J. Immunol. 129, 1627–1631 (1982).

49. Voisin, T. et al. The CysLT2R receptor mediates leukotriene C4-driven acute and chronic itch. Proc. Natl. Acad. Sci. U. S. A. 118, e2022087118 (2021).

50. Wang, F. et al. A basophil-neuronal axis promotes itch. Cell 184, 422-440.e17 (2021).

51. Prescott, S. L., Umans, B. D., Williams, E. K., Brust, R. D. & Liberles, S. D. An Airway Protection Program Revealed by Sweeping Genetic Control of Vagal Afferents. Cell 181, 574-589.e14 (2020).

52. Taylor-Clark, T. E. et al. Prostaglandin-induced activation of nociceptive neurons via direct interaction with transient receptor potential A1 (TRPA1). Mol. Pharmacol. 73, 274–281 (2008).

53. Zhang, S. et al. Role of prostaglandin D2 in mast cell activation-induced sensitization of esophageal vagal afferents. Am. J. Physiol. Gastrointest. Liver Physiol. 304, G908–916 (2013).

54. Xu, H. et al. Transcriptional Atlas of Intestinal Immune Cells Reveals that Neuropeptide α-CGRP Modulates Group 2 Innate Lymphoid Cell Responses. Immunity 51, 696-708.e9 (2019).

55. Roper, S. D. TRPs in Taste and Chemesthesis. Handb. Exp. Pharmacol. 223, 827–871 (2014).

56. Zhang, C. et al. Area Postrema Cell Types that Mediate Nausea-Associated Behaviors. Neuron 109, 461-472.e5 (2021).

57. Worth, A. A. et al. The cytokine GDF15 signals through a population of brainstem cholecystokinin neurons to mediate anorectic signalling. eLife 9, e55164 (2020).

58. Tsai, V. W.-W. et al. The anorectic actions of the TGFβ cytokine MIC-1/GDF15 require an intact brainstem area postrema and nucleus of the solitary tract. PloS One 9, e100370 (2014).

59. Patel, S. et al. GDF15 Provides an Endocrine Signal of Nutritional Stress in Mice and Humans. Cell Metab. 29, 707-718.e8 (2019).

60. Eisenstein, A. et al. Activation of the transcription factor NRF2 mediates the anti-inflammatory properties of a subset of over-the-counter and prescription NSAIDs. Immunity S1074-7613(22)00186–8 (2022) doi:10.1016/j.immuni.2022.04.015.

61. Rankin, L. C. & Artis, D. Beyond Host Defense: Emerging Functions of the Immune System in Regulating Complex Tissue Physiology. Cell 173, 554–567 (2018).

62. Tracey, K. J. Reflex control of immunity. Nat. Rev. Immunol. 9, 418–428 (2009).

63. Koren, T. et al. Insular cortex neurons encode and retrieve specific immune responses. Cell 184, 5902-5915.e17 (2021).

64. Yu, C. Targeted Deletion of a High-Affinity GATA-binding Site in the GATA-1 Promoter Leads to Selective Loss of the Eosinophil Lineage In Vivo. J. Exp. Med. 195, 1387–1395 (2002).

65. Noben-Trauth, N. et al. An interleukin 4 (IL-4)-independent pathway for CD4+ T cell IL-4 production is revealed in IL-4 receptor-deficient mice. Proc. Natl. Acad. Sci. 94, 10838– 10843 (1997).

66. Florsheim, E. et al. Integrated Innate Mechanisms Involved in Airway Allergic Inflammation to the Serine Protease Subtilisin. J. Immunol. 194, 4621–4630 (2015).

67. Luan, H. H. et al. GDF15 Is an Inflammation-Induced Central Mediator of Tissue Tolerance. Cell 178, 1231-1244.e11 (2019).

68. Schindelin, J. et al. Fiji: an open-source platform for biological-image analysis. Nat. Methods 9, 676–682 (2012).

69. Spencer, S. P. et al. Adaptation of Innate Lymphoid Cells to a Micronutrient Deficiency Promotes Type 2 Barrier Immunity. Science 343, 432–437 (2014).

70. Sullivan, Z. A. et al. γδ T cells regulate the intestinal response to nutrient sensing. Science 371, eaba8310 (2021).

